# Analysis of single-cell gene pair coexpression landscapes by stochastic kinetic modeling reveals gene-pair interactions in development

**DOI:** 10.1101/815878

**Authors:** Cameron P. Gallivan, Honglei Ren, Elizabeth L. Read

**Affiliations:** Dept. of Chemical & Biomolecular Engineering, University of California, Irvine, CA, USA; NSF-Simons Center for Multiscale Cell Fate, University of California, Irvine, CA, USA; Mathematical and Computational Systems Biology Graduate Program, University of California, Irvine, CA, USA

**Keywords:** stochastic modeling, gene expression noise, gene regulatory networks, single-cell data, scRNA-seq

## Abstract

Single-cell transcriptomics is advancing discovery of the molecular determinants of cell identity, while spurring development of novel data analysis methods. Stochastic mathematical models of gene regulatory networks help unravel the dynamic, molecular mechanisms underlying cell-to-cell heterogeneity, and can thus aid interpretation of heterogeneous cell-states revealed by single-cell measurements. However, integrating stochastic gene network models with single cell data is challenging. Here, we present a method for analyzing single-cell gene-pair coexpression patterns, based on biophysical models of stochastic gene expression and interaction dynamics. We first developed a high-computational-throughput approach to stochastic modeling of gene-pair coexpression landscapes, based on numerical solution of gene network Master Equations. We then comprehensively catalogued coexpression patterns arising from tens of thousands of gene-gene interaction models with different biochemical kinetic parameters and regulatory interactions. From the computed landscapes, we obtain a low-dimensional “shape-space” describing distinct types of coexpression patterns. We applied the theoretical results to analysis of published single cell RNA sequencing data and uncovered complex dynamics of coexpression among gene pairs during embryonic development. Our approach provides a generalizable framework for inferring evolution of gene-gene interactions during critical cell-state transitions.

## 1 INTRODUCTION

In recent years, single-cell-resolution measurements have revealed unprecedented levels of cell-to-cell heterogeneity within tissues. The discovery of this ever-present heterogeneity is driving a more nuanced view of cell phenotype, wherein cells exist along a continuum of cell-states, rather than conforming to discrete classifications. The comprehensive view of diverse cell states revealed by single cell measurements is also affording new opportunities to discover molecular regulators of cell phenotype and dynamics of lineage commitment (Trapnell et al. (2014); Olsson et al. (2016); Briggs et al. (2018)). For example, single cell transcriptomics have revealed the widespread nature of *multilineage priming* (MLP), a phenomenon wherein individual, multipotent cells exhibit “promiscuous” coexpression of genes associated with distinct lineages prior to commitment (Nimmo et al. (2015)). In principle, mathematical modeling of gene regulatory network dynamics can provide a theoretical foundation for understanding cell heterogeneity and gene expression dynamics, by quantitatively linking molecular-level regulatory mechanisms with observed cell states. However, due to the molecular complexity of gene regulatory mechanisms, it remains challenging to integrate such models with single-cell data.

Mathematical models of gene regulatory network dynamics can account for (and at least partially reproduce) observed cellular heterogeneity in two primary ways. First, gene network models are multi-stable dynamical systems, meaning a given network has the potential to reach multiple stable states of gene expression. These states arise from the dynamic interplay of activation, inhibition, feedback, and nonlinearity (Kauffman (1969); MacArthur et al. (2009); Huang (2012)). Second, some mathematical models inherently treat cellular noise. This noise, or stochasticity, is modeled in various ways depending on assumptions about the source (Peccoud and Ycart (1995); Arkin et al. (1998); Kepler and Elston (2001); Swain et al. (2002)). Discrete, stochastic models of gene regulation, which track discrete molecular entities, regulatory-protein binding kinetics, and binding states of promoters controlling gene activity, have formed the basis of biophysical theories of gene expression noise due to so-called *intrinsic* molecular noise (Peccoud and Ycart (1995); Thattai and van Oudenaarden (2001); Kepler and Elston (2001); Pedraza and Paulsson (2008)). Such stochastic gene-regulation mechanisms have also been incorporated into larger regulatory network models using the formalism of stochastic biochemical reaction networks, and have been utilized to explore how molecular fluctuations can cause heterogeneity within phenotype-states and promote stochastic transitions between phenotypes (Feng and Wang (2012); Sasai et al. (2013); Zhang and Wolynes (2014); Tse et al. (2015)).

The quantitative *landscape* of cellular states is another concept that is increasingly utilized to describe cellular heterogeneity. Broadly, the cellular potential landscape (first conceptualized by Waddington (Waddington (2014); Wang et al. (2011); Huang (2012)) is a function in high-dimensional space (over many molecular observables, typically expression levels of different genes), that quantifies the stability of a given cell-state. In analogy to potential energy (gravitational, chemical, electric, etc.), cell states of higher potential are less stable than those of lower potential. The landscape concept inherently accounts for cellular heterogeneity, since it holds that a continuum of states is theoretically accessible to the cell, with low-potential states (in “valleys”) more likely to be observed than high-potential states. The landscape is a rigorously defined function derived from the dynamics of the underlying gene network model, according to some choice of mathematical formalism (Wang et al. (2011); Bhattacharya et al. (2011); Huang (2012); Zhou et al. (2016)). For stochastic gene network models that inherently treat noise, the landscape is directly obtained from the computed probability distribution over cell-states (Cao and Liang (2008); Micheelsen et al. (2010); Feng and Wang (2012); Tse et al. (2015)).

Stochastic modeling of gene network dynamics has been employed in various forms for analysis of single cell measurements. For example, application of noisy dynamical systems theory has shed light on cell-state transitions (Mojtahedi et al. (2016); Jin et al. (2018); Lin et al. (2018)). Stochastic simulations of gene network dynamics have been used to develop and/or benchmark tools for network reconstruction (Schaffter et al. (2011); Dibaeinia and Sinha (2019); Bonnaffoux et al. (2019)) Stochastic model-aided analysis of single-cell measurements has been demonstrated to yield insights on gene regulatory mechanisms (Munsky et al. (2018)). However, few existing analysis methods utilize discrete-molecule, stochastic models, which fully account for intrinsic gene expression noise and its impact on cell-state, to aid in the interpretation of noisy distributions recovered from single cell RNA sequencing data. There exists an opportunity to link such biophysical, stochastic models, which reproduce intrinsic noise and cell heterogeneity *in silico*, to single cell datasets that characterize cell heterogeneity *in vivo*. In particular, the landscape of heterogeneous cell-states computed from discrete stochastic models can be directly compared to single-cell measurements.

In this work, we present a method for analyzing single-cell gene pair coexpression patterns that is founded on biophysical theory of stochastic gene networks. In our approach, the key object linking the models to the data is the gene-pair coexpression landscape, which is derived directly from the bivariate distribution of expression states, and which is computed from a stochastic model or extracted from single cell measurements. The rationale underlying the method is two-fold: (1) information on gene-gene interactions can be inferred from the distinctive characteristics of noise in single-cell data (i.e., from the “shape” of the landscape); (2) existing analysis techniques are relatively insensitive to landscape shape. We first comprehensively compute and classify the landscapes produced by a family of ~40,000 stochastic two-gene regulatory network models. We then use the model-derived classification to analyze published data from vertebrate development. In so doing, we uncover both expected and novel patterns of coexpression in development. While our analysis here is proof-of-principle, and limited to two-gene interactions, the conceptual framework could be expanded to include multi-body gene interactions in the future.

## 2 METHODS

### 2.1 Discrete, Stochastic Models of Two-Gene Regulatory Networks

We first developed a family of stochastic models of gene-gene interactions (see Fig. 1 for model schematic), which is based on previously published models (Feng and Wang (2012); Zhang and Wolynes (2014)). We label two genes *X* and *Y*. Each gene encodes a protein, which acts as a transcription factor (TF) that potentially regulates its own expression as well as that of the other gene. Each gene has a promoter (or more generally, regulatory regions of DNA) that can be bound by any combination of its own expressed protein and/or the other gene’s expressed protein. The promoter states are thus labeled as: *X*_00_ (neither transcription factor is bound to X’s promoter), *X*_0*x*_ (X’s own protein is bound, resulting in auto-regulation of gene expression), *X*_*y*0_ (Y’s protein is bound to X’s promoter, resulting in cross-regulation), *X*_*yx*_ (both proteins are bound to X’s promoter, resulting in combinatorial regulation). (The promoter states for gene *Y* are defined in a symmetric manner.) The regulatory effect of each promoter state (i.e., the effect of having none, one, or both proteins bound on the gene’s expression) is accounted by the transcription rate *g*_*ij*_ corresponding to each possible promoter state: e.g., when gene *X*’s promoter is unbound, it transcribes at rate 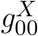. Binding of *Y*’s protein changes the transcription rate to 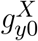, which may be lower, higher, or the same, depending on whether the effect of *Y* on *X* is assumed to be repressing, activating, or not impacting. (All other transcription rates for each promoter state and for gene *Y* are defined similarly.) The model involves three classes of reactions: mRNA synthesis, mRNA degradation, and promoter-state-change reactions. mRNA synthesis reactions are given by:

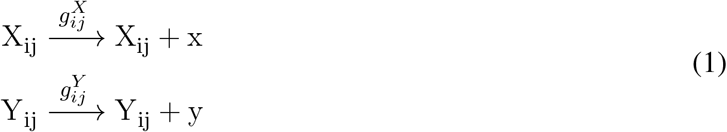

where *x* and *y* denote mRNA transcripts which will be translated into the transcription factors encoded by genes *X* and *Y*, respectively. mRNA degradation reactions are given by:

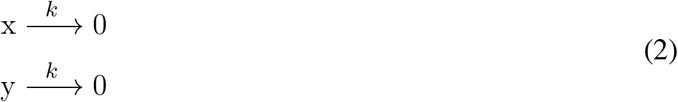

**Figure 1.**
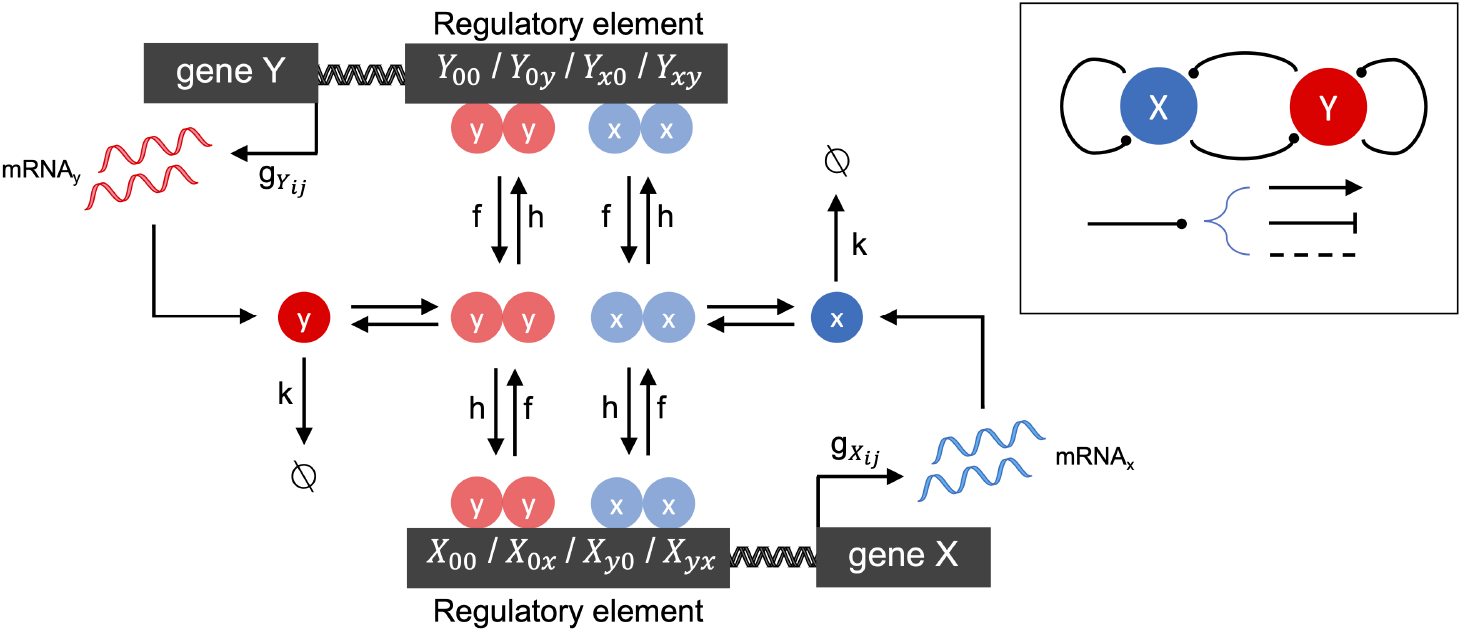
Schematic of the two-gene regulatory network model. The overall network motif is variable (see Inset), encoding a symmetric combination of repression (flat arrow-head), activation (pointed arrow-head) or no-impact (dashed line), mutually between the two genes labeled *X* and *Y*, and by each gene on itself (see Methods for details). The stochastic reaction kinetic model includes rate constants for mRNA synthesis (*g*_*ij*_), mRNA degradation (*k*), and regulatory element state-changes due to transcription factor binding (*h*) and unbinding (*f*). Cooperative effects are included by the assumption that transcription factors bind as homodimers.

Promoter-state-change reactions are given by, e.g.:

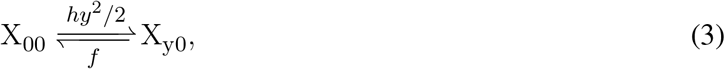

which represents the change of promoter-state (and corresponding regulatory impact) on gene *X* when *Y*’s transcription factor binds (forward reaction) or unbinds (reverse reaction). All other promoter-state-change reactions for *X* and *Y* are defined similarly. The changes of promoter state occur with forward rates *hy*^2^/2 or *hx*^2^/2 (when the change of state occurs due to binding of transcription factor from gene *Y* or *X*, respectively) and *f* (when the change of state occurs due to an unbinding event). The model tracks copy numbers of individual mRNA molecules in the cell, to enable direct comparison with single cell transcriptomic data, but translation of mRNA into protein is not explicitly accounted for. Instead, transcription factor (protein) levels are assumed to be linearly proportional to mRNA, and this proportionality constant is subsumed into the binding rate *h*. The quadratic dependence of the forward binding rates on *x* or *y* arises from the assumption that homodimeric transcription factors regulate gene expression, which is a general and convenient way to include cooperativity in the model.

We assign rate constants to intracellular processes that are in line with experimental estimates from vertebrates, where possible (see Table 1). (For full details of model reactions and parameter derivations, see Supplement). Rates of mRNA synthesis and degradation are relatively well characterized, though they vary considerably for different transcripts (Schwanhäusser et al. (2011)). Rates of promoter-state-change are less well-defined, since promoter-state-changes that ultimately impact gene expression may be attributed to a variety of molecular processes, including: (a) relatively fast processes of TF binding or unbinding from DNA (b) relatively slow chromatin remodeling processes that may be initiated or facilitated by TF binding, require multiple steps and cooperative interactions, and are generally poorly understood. In our models, to account for this range of possible mechanisms, we consider a wide range of parameter values *h, f* for promoter-state-changes. (The significance of these fast and slow regimes, termed the *adiabatic* and *nonadiabatic* regimes, respectively, to cell-state stability has been studied previously by stochastic modeling (Sasai and Wolynes (2003); Feng and Wang (2012); Sasai et al. (2013); Zhang and Wolynes (2014))). We here define the “fast” regime as determined by measured parameter values of protein binding/unbinding DNA (e.g., from Geertz et al. (2012)), occurring with timescales of minutes, seconds, or faster. We define the “slow” regime more broadly as any epigenetic/chromatin changes occurring on timescales of hours, days, or longer. For example, in mammalian cells, changes of chromatin state during cell-fate specification were estimated to be on the order of several days (Hathaway et al. (2012); Mariani et al. (2010)), while theoretical studies predicted timescales on the order of the cell cycle time (i.e., hours to days, Sasai et al. (2013)).

**Table 1.** Rate Parameters used in gene regulatory network models. Parameter values are derived from experimental measurements in vertebrates, where possible. See Methods text for details. *Measured rates of mRNA synthesis varied, with a median of 2/hr Schwanhäusser et al. (2011)). We use lower values (within experimental range) to roughly match observed counts in scRNA-seq data, which may be lower than expected because of dropouts or other technical issues. ^‡^Corresponds to mRNA half-life of 3.5 hours, which is well within experimentally measured values but shorter than the median value of 9 hours, assuming that transcriptional regulators have shorter-than-average half-lives in the cell. *†* Promoter state change rates *f* and *k* are reported in fast and slow regimes. Fast promoter state changes are assumed to occur due to TF-DNA unbinding or binding events, with rate parameters chosen based on values reported in Geertz et al. (2012) (see Supplement for details on parameter derivation and unit conversion). Slow promoter state changes are thought to involve collective changes in epigenetic marks and rearrangement of chromatin.

| Rate Constant | Symbol | Units | Value | Comments/Source |
| --- | --- | --- | --- | --- |
| mRNA synthesis (not repressed) | $g_{hi}$ | mRNA/hr | 0.8 - 1.4* | Schwanhäusser et al. (2011) |
| mRNA synthesis (repressed) | $g_{lo}$ | mRNA/hr | 0.001 | see text |
| mRNA degradation | $k$ | /hr | $0.2^{\ddagger}$ | Schwanhäusser et al. (2011) |
| Promoter state change (unbinding) | $f^{\dagger}$ | /hr | (fast) $10 - 10^5$<br>(slow) $10^{-6} - 10$ | Geertz et al. (2012)<br>see text |
| Promoter state change (binding) | $h^{\dagger}$ | $\text{hr}^{-1} \text{mRNA}^{-2}$ | (fast) $10 - 500$<br>(slow) $10^{-6} - 10$ | Geertz et al. (2012)<br>see text |

We define two types of model systems. The **Mutual Inhibition/Self-Activation (MISA)** model encodes a common network motif that is understood to control a variety of cell fate decisions (Graf and Enver (2009); Huang (2013)) and has been extensively studied by mathematical modeling (Huang et al. (2007); Feng and Wang (2012); Chu et al. (2017)). In contrast, the **Two-Gene Flex** model flexibly encodes a variety of regulatory interactions, as described below.

#### 2.1.1 Mutual Inhibition/Self-Activation Model

In all models, promoter activity is assumed to be either high (transcription rate *g*_*hi*_) or low (*g*_*lo*_) (giving a relatively fast or slow rate of mRNA synthesis, respectively). To encode MISA regulatory logic, mRNA synthesis rates for each promoter state are 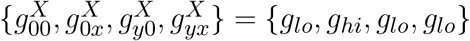. Transcription rates for gene *Y* are defined symmetrically, 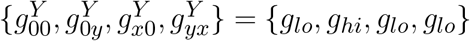. The high rate corresponds to maximal activity, whereas the low rate is effectively off (but is non-zero to allow for some leakiness in the promoter). Thus, binding of the self-TF turns the gene on, but subsequent binding of the other TF turns the gene off. The relative strengths and kinetics of the activating (self-regulatory) and repressing (cross-regulatory) interactions are encoded in the rates of binding/unbinding of regulators. Autoregulatory binding and unbinding rates (symmetric on both genes) are denoted by *h*_*a*_ and *f*_*a*_, respectively. Cross-regulatory rates are denoted by *h*_*r*_ and *f*_*r*_. The model is thus fully specified by 7 parameters: {*g*_*lo*_, *g*_*hi*_, *k, h*_*a*_, *f*_*a*_, *h*_*r*_, *f*_*r*_}. We computed landscapes for ~22,000 unique parameter combinations for the MISA regulatory logic (see Table 1 for parameter value ranges). We studied only symmetric network motifs, but asymmetry between the genes is accounted for by allowing the “on” transcription rate *g*_*hi*_ to be asymmetric between the two genes (in case of asymmetry in *g*_*hi*_, the model is specified by eight parameters).

#### 2.1.2 Two-Gene Flex Model

The Two-Gene Flex model is identical to MISA in all ways except the regulatory logic. Instead of the transcription rates being {*g*_*lo*_, *g*_*hi*_, *g*_*lo*_, *g*_*lo*_}, all 16 logical combinations of four promoter states and two activity-levels are included. Within these combinations, various behavior is encoded including self-activation, self-repression, mutual activation, mutual repression, no interaction (self- or cross-), and dual-effects (where a TF has a distinct effect whether bound alone or in combination with the other). Note that the MISA logic is contained within these 16 combinations. Note also that the promoter states for *X* and *Y* are always defined symmetrically, i.e., only symmetric motifs are included. We computed landscapes for ~34,000 unique parameter combinations for the Two-Gene Flex Model (including all network motif variants). Our aim with the Two-gene Flex model was to comprehensively encode all possible logical combinations within the constraints of the symmetric two-gene model. Note that these combinations encompass several cis-regulatory motifs that have been described previously. For example, {*g*_00_, *g*_0*y*_, *g*_*x*0_, *g*_*yx*_} = {*g*_*hi*_, *g*_*lo*_, *g*_*hi*_, *g*_*lo*_} corresponds to a “simple repressor” motif where Y is the repressor, and {*g*_00_, *g*_0*y*_, *g*_*x*0_, *g*_*yx*_ = *g*_*hi*_, *g*_*lo*_, *g*_*lo*_, *g*_*lo*_} corresponds to a “dual repressor” motif (Bintu et al. (2005)). Our Two-Gene Flex model also encompasses various biologically-inspired logic gates for combinatorial cis-regulation studied previously (Zhang et al. (2009)).

### 2.2 Mathematical Framework: Chemical Master Equation

#### 2.2.1 Chemical Master Equation

Stochastic dynamics for the above-described network motifs are modeled by a Chemical Master Equation (CME) (alternatively known as a discrete space, continuous time Markov Chain). The instantaneous state of the system is given by the vector **n**, which enumerates the mRNA copy numbers and promoter-states of both genes, i.e., **n**= [*n*_*x*_, *n*_*y*_, *X*_*ij*_, *Y*_*ij*_], where *n*_*x*_ is the mRNA copy number for gene *X*, *X*_*ij*_ is the promoter state for gene *X*, and so on. The CME gives the probability for the system to exist in a given state at a given time, **p**(**n**, *t*). The CME can be written in vector-matrix form as a linear system

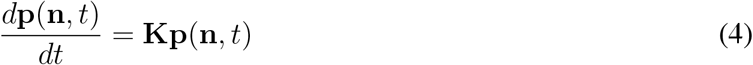

where **K** is the reaction rate-matrix. Each off-diagonal element *K*_*lm*_ gives the rate of transitioning from state *m* to *l* (non-zero values correspond to allowed state transitions with rates according to reactions 1-3 above), while the diagonal elements are the summed rates for exiting each state, *K*_*ll*_ = −∑_*m*≠*l*_ *K*_*ml*_. Transition rates are computed according to standard stochastic chemical kinetic rate laws (Gillespie (1977)). If both types of mRNA are assumed to exist in the cell in copy numbers that never exceed *M* − 1, then the total size of the enumerated space including all possible states is *N* = *M* × *M* × 4 × 4 (note that the total number of mRNA copy number states includes the state of 0 copies, thus *n*_*x*_, *n*_*y*_ ∈ {0, 1, …*M* − 1}). The assumption that mRNAs never exceed *M* − 1 is equivalent to assuming reflective boundary conditions on the enumerated state-space. That is, it assumes the propensity of reactions that lead to mRNA numbers exceeding *M* − 1 is 0. This assumption is justified when *M* is chosen to be sufficiently large compared to *g*/*k* (Chu et al. (2017)). We confirmed that the probability of mRNAs exceeding *M* −1 for our parameter values is negligible (Supplement, Section 2.2) and we further confirmed that increasing *M* (from 21, the value used in calculations throughout the manuscript, to 36) had negligible impact on quasipotential landscape shape and all subsequent analysis of single cell RNA sequencing (scRNA-seq) data (Fig. S2). Note that an algorithm has been published recently that provides rigorous error bounds on steady-state solutions to the CME (Gupta et al. (2017)) though we do not make use of the algorithm here.

#### 2.2.2 Computing Gene Pair Coexpression Landscapes

The complete steady state probability to find a cell in state **n** is given by the vector ***π*** (**n**) = **p**(**n**, *t* → ∞), which is obtained from Eq. 4 using eigenvalue routines in numpy and scipy (van der Walt et al. (2011)) (McKinney (2010)). Each individual model requires solution of an *N*-state system, where *N* is 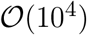 (e.g., assuming the probability to have mRNA exceed 25 is negligible, then *N* = 10, 816). Efficient computation of the landscapes over tens of thousands of model variants/parameter combinations was achieved using routines compiled with the numba library (Lam et al. (2015)) and parallelization using Python’s multiprocessing library to distribute the workload across the available cores.

To mimic experimental scRNA-seq data, the probability is projected onto the mRNA subspace by summation over all promoter state combinations. We hereon define the gene pair coexpression landscape as the steady-state probability to find a cell with mRNA count numbers (*n*_*x*_, *n*_*y*_). More precisely, the *probability landscape* is the vector ***π*** with each element *π*_*i*_ giving the steady-state probability for the cell to be found in state *i* with the combination of mRNA counts (*n*_*x*_, *n*_*y*_) from genes *X* and *Y*, and *i* ∈ 1, …, *M*^2^. Alternatively, the *quasipotential landscape* is log-transformed, given by the vector ***ϕ*** where *ϕ*_*i*_ = −ln(*π*_*i*_).

### 2.3 scRNA-seq Data Acquisition, and Landscape Estimation

Experimental data is obtained from the published scRNA-seq measurements of Briggs et al. (2018). The dataset “Corrected combined.annotated counts.tsv” was used which provides the normalized transcriptome profiles for *Xenopus tropicalis* at single cell resolution for ten different stages of embryonic development, with labeled cell types and parent cell types. We analyzed 1380 gene pairs, which were identified as putative MLP pairs in Briggs et al. (2018), based on their estimated changes in coexpression over the course of development. Gene pairs were identified by their developmental stage and lineage branch point in which coexpression was maximal. Cell types from other stages were then included in the lineage if they were a parent (preceding in development) cell type or daughter (descendant later in development) cell type. After selecting the desired gene pair and cell/tissue/cluster type of interest, gene pair counts were combined and summed resulting in ten gene pair landscapes, one for each stage of development, in cells of the relevant lineage.

To directly compare computed coexpression landscapes with experimental data, we extracted cell count matrices for each gene pair, and where necessary, truncated to mRNA count numbers ≤ *M* − 1 (truncation eliminated less than 0.5% of cells in the data, across all gene pairs and cell stages). This produces an *M × M* (including zeros) count matrix that serves as a sampled estimator of the steady-state distribution, 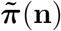, of the same size as computed landscapes. In order to compute the sampled quasipotential landscape, we use 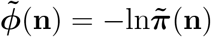, after replacing the not-observed count-combinations with a low but non-zero estimate of these probabilities (since log of zero is undefined). We use a general estimate of 1E-6 for non-observed counts, both because it is in line with the predictions of the theoretical models for the low probability edges of the distributions, and because it is less than the lowest estimable probability (i.e., observation of one cell in a given matrix position, given total cell counts on the order of 10^5^, would correspond to an estimated probability of 1E-5).

### 2.4 Dimensionality Reduction for Landscape Shape-space

We apply Principal Component Analysis (PCA) to the theoretically computed landscapes over the model sets to achieve a reduced-dimension description of landscape shape. All PCA training and dimensionality reduction was performed using the decomposition module of the python package scikit learn. Each unique model is treated as a replicate and the steady-state probability *π*_*i*_ (or alternatively, quasipotential *ϕ*_*i*_) of each of the *M* × *M* possible mRNA copy-number states (*n*_*x*_, *n*_*y*_) is treated as a feature.

The principal components obtained from the model set were then used to fit the experimental data, where each landscape from each gene-pair/stage is a replicate. Note that in our application, we have opted to use a “theory-driven” analysis of landscape shape-space, where the PCA training set consists of theoretically computed probability (or quasipotential) landscapes. The experiment-derived landscapes are then projected into this theory-driven shape-space, which enables linking of experimentally measured gene-pair landscapes with possible model logic/parameter combinations that could produce observed landscape shapes. Alternatively, a “data-driven” analysis is possible, wherein the PCA training set consists of experiment-derived landscapes. Such an analysis makes no connection between theoretical models and experimental data, but can still be useful in revealing shape-features present in experimental data. We show results from data-driven analysis in Supplement section 2.4 and Fig. S5.

### 2.5 Clustering of Developmental Landscape-Shape Trajectories

By viewing the time-ordered coexpression landscapes of a given gene pair in PCA space, termed “landscape-shape trajectories”, one can gain insight into the genes’ roles in development. The trajectories were hierarchically clustered based on their geometric distance in PCA space. More specifically, the *fcluster* method in scikit-learn package was used in hierarchical clustering (McKinney (2010)), and the geometric distance between trajectories A and B were defined as the sum of the pair-wised Euclidean distance between two corresponding stages, i.e.

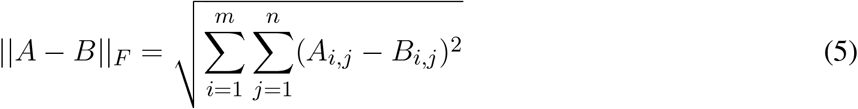

where ‖·‖_*F*_ is the Frobenius norm, *A* and *B* are two trajectories represented by *m* by *n* matrices, *m* is the number of developmental stages in single cell data, *n* is the number of PCA components used in clustering.

## 3 RESULTS

### 3.1 Stochastic two-gene network models show a variety of coexpression landscape shapes, distinguishable by Principal Component Analysis

Our modeling framework enabled efficient computation of coexpression landscapes resulting from discrete, stochastic gene network models. This in turn enabled us to compute landscapes for tens of thousands of parameter sets, encompassing both various relative strengths and kinetics of regulatory interactions, as well as different schemes of regulatory logic among the two genes (see Methods). This approach afforded a comprehensive view of theoretically predicted landscape shapes resulting from gene-gene interactions (within the assumptions of the current model system).

We applied Principal Component Analysis to the computed probability landscapes for Two-Gene Flex, in order to find a low-dimensional description of their shapes (Fig. 2). The first two PCA components encompass 98% percent of total covariance, and all models fall within a triangular region of this 2D subspace. The vertices of the triangle correspond generally to landscapes with: (1) very low expression of both genes (i.e., transcript levels of *X/Y* are *lo/lo*, Fig. 2E), (2) high simultaneous expression of both genes (*hi/hi*, Fig. 2C), and (3) expression of only one gene at a time (*hi/lo* and *lo/hi*, Fig. 2A). Landscapes located away from the vertices are thus well-described by some linear combination of these three shapes, consistent with PCA, and supported by visual inspection. In all, the results reveal that two-gene interaction motifs can encode a wide variety of patterns of coexpression, including mixtures of all combinations of *lo/lo*, *hi/hi* and *lo/hi*, *hi/lo* phenotypes (e.g,. Fig. 2B). At the same time, this variety of shapes is well-described by a small number of principal components (which form a basis for what we term the “shape-space”), and we hereon use the magnitudes along these components as measures of landscape shape.

**Figure 2.**
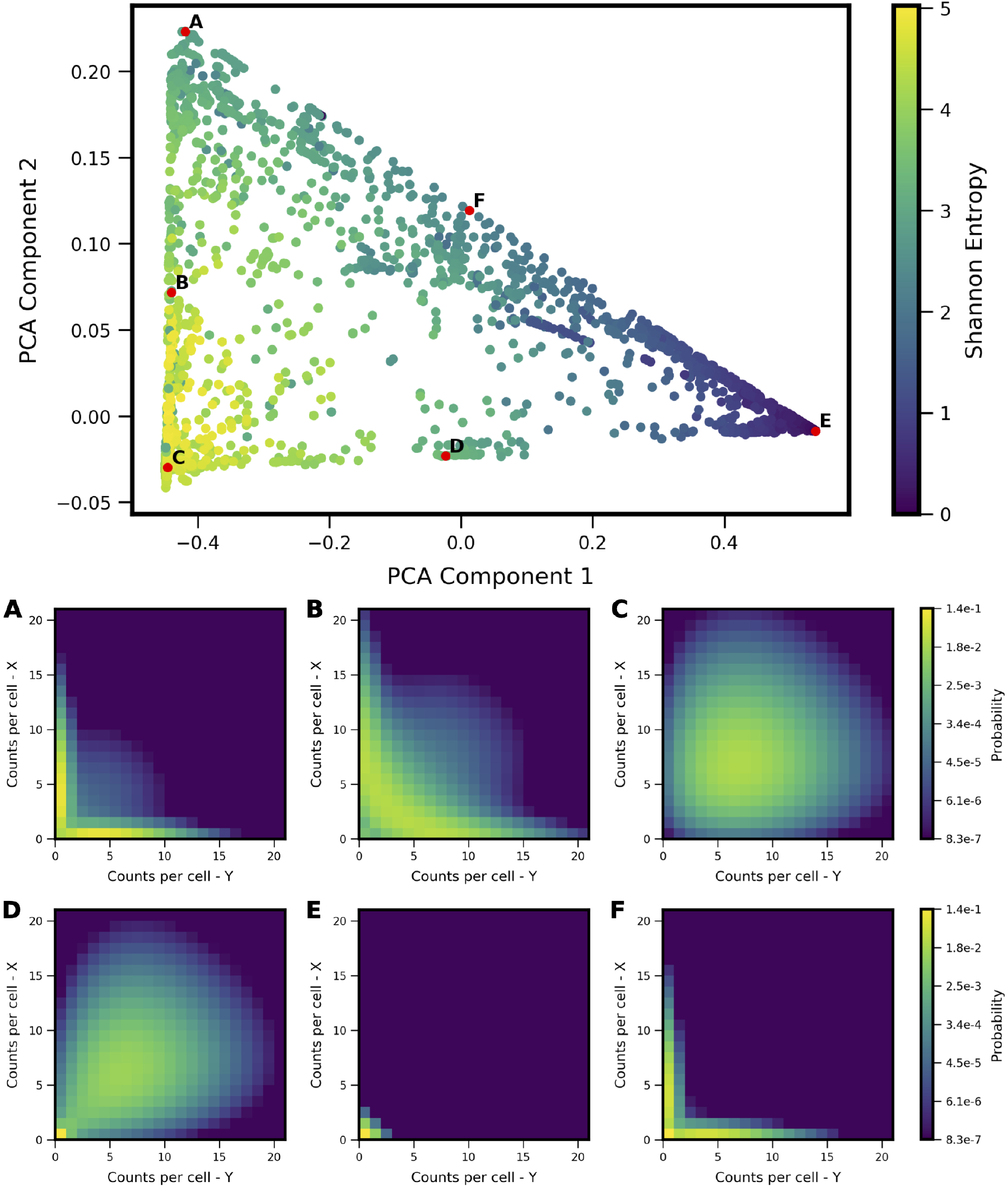
Shape-space of simulated Two-Gene Flex coexpression landscapes analyzed by PCA. Co-expression landscapes were computed for 34,097 unique two-gene stochastic network models with varying regulatory interactions and kinetic rate parameters (see Model schematic in Fig. 1). (Top) All model landscapes projected onto the first two principal components. Each dot corresponds to one model, colored by the model’s Shannon Entropy. (Bottom) Representative quasipotential landscapes *ϕ*(*n*) (see Methods) of individual models from different regions of PCA component-space. Color of each discrete grid space in {*x, y*} corresponds to computed probability (in log-scale) to find a single cell with the corresponding numbers of {*x, y*} transcripts.

### 3.2 Shape measures of coexpression landscapes distinguish different types of mutual gene-gene interactions

We sought to understand how different regulatory motifs contributed to landscape shape. Projecting the landscapes arising from each network motif separately revealed distinctive patterns (i.e., occupying distinct, but overlapping, regions of the PCA triangle) (approximately 2,000 landscapes were computed for each network motif, i.e., ~2,000 models that share the regulatory logic but have different kinetic parameters). We grouped all motifs according to their region of occupancy within the PCA triangle, and discovered logical consistency among the groups (see Fig. 3). For example, all motifs with some type of mutual activation were found to co-occupy a region of PCA shape-space in the lower part of the triangle (3A). This result is consistent with the intuition that motifs with mutual activation cannot produce the apparent bistability seen in landscapes at the *hi/lo*-*lo/hi* vertex of the triangle. The other three motif groupings include motifs with some type of mutual repression, motifs with no inter-gene interactions, and incoherent motifs with dual-interactions (when the regulator bound by itself has the opposite effect of the regulator bound in combination with the other TF). Note that two of the sixteen logical combinations of promoter binding-states in the Two-Gene Flex models are not included here, since they effectively encode no gene-gene interactions (the “always on” or “always off” logic, {*g*_*hi*_, *g*_*hi*_, *g*_*hi*_, *g*_*hi*_} or {*g*_*lo*_, *g*_*lo*_, *g*_*lo*_, *g*_*lo*_}). Note that here we assess all kinetic parameter combinations associated to one regulatory motif; these parameters tune the strength of different interactions. As such, the analysis of Fig. 3 assumes fixed network topologies but variable weights on network edges, accounting for the overlap between different motifs. These results indicate that landscape shape can to some extent be used to distinguish regulatory interactions between pairs of genes, despite variable and/or unknown kinetics governing the interactions.

**Figure 3.**
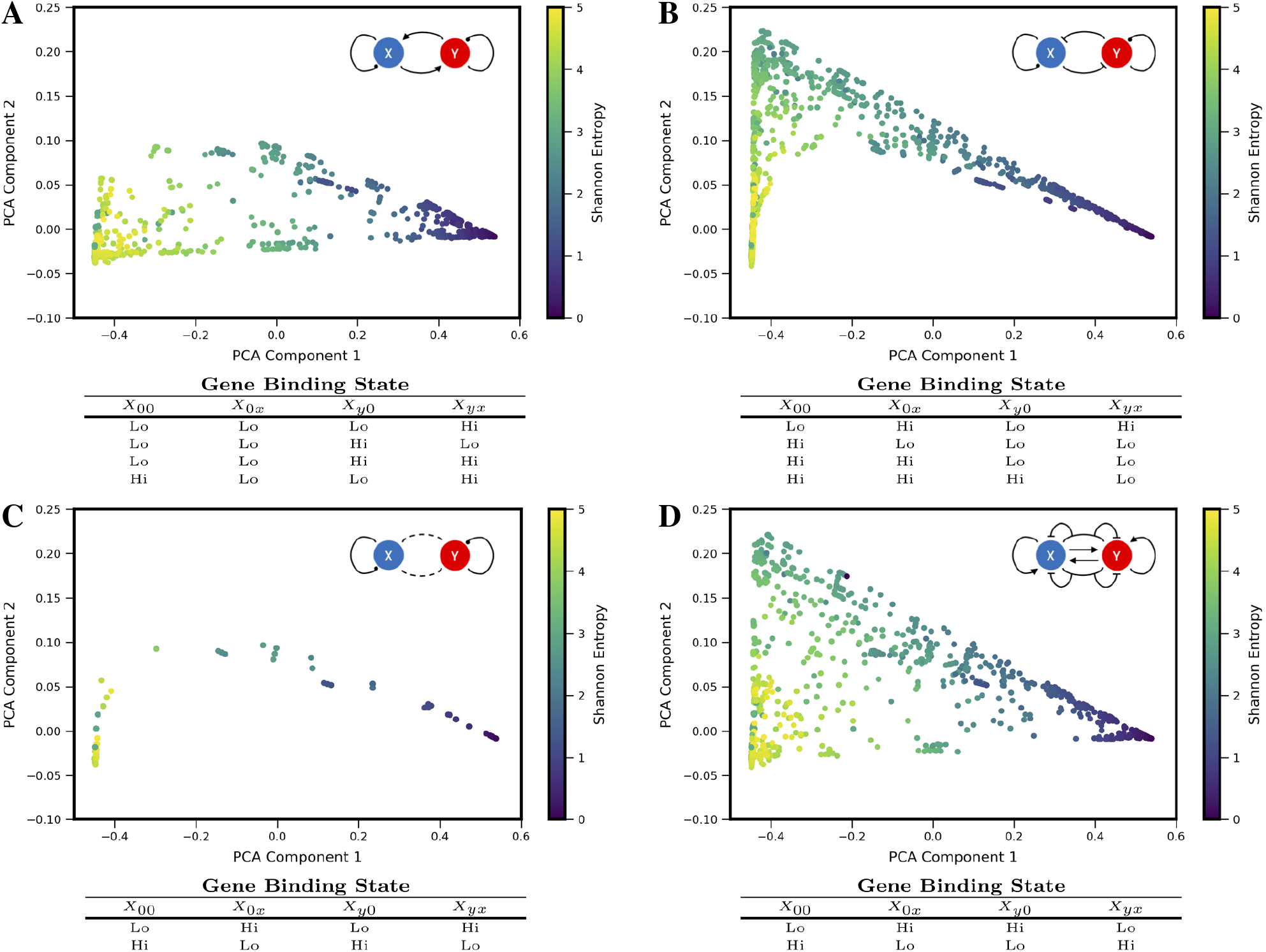
Coexpression landscapes computed from the Two-Gene Flex models show distinctive shapes that depend on the regulatory logic of gene-gene interactions. The Two-Gene Flex model encodes 16 logical combinations (2^4^) of gene-gene interactions, corresponding to four possible promoter-binding states and two possible levels of transcription activity (low and high). These 16 model variants can be grouped into motif classes: (**A**) Models with mutual activation. (**B**) Models with mutual repression. (**C**) No mutual gene-gene interactions. (**D**) “Incoherent” models, where the combinatorial-binding state has the opposite behavior of both of the singly-bound states (see text). Within each motif class, different kinetic parameters serve to modify the relative strength of interactions (i.e., different weights on the edges). Each motif class occupies a distinct, but overlapping, region of the shape-space (with the exception of the Incoherent motif, which can reach all areas of the shape-space).

### 3.3 Commonly used pairwise metrics are relatively insensitive to coexpression landscape shape

In order to analyze how previously-applied measures of gene-gene interactions align with landscape shape, we computed a set of metrics for each model landscape and visualized the resultant values projected onto the PCA subspace. We chose four metrics: Shannon Entropy, Pearson Correlation Coefficient, Mutual Information, and a Coexpression Index (see Fig. 4, note Shannon Entropy is visualized also in Figs. 2 and 3). The first three of these are obtained directly from the computed bivariate probability distributions according to standard definitions; the Coexpression Index has been used previously (Briggs et al. (2018)) and is given by the conditional probability to find cells with non-zero counts of both mRNA *x* and *y* (conditioned on the cells having non-zero counts of at least one of genes *X* or *Y*). Here, for a given model *j*, we derive this metric from the probability landscape *π* over count-states *i* by:

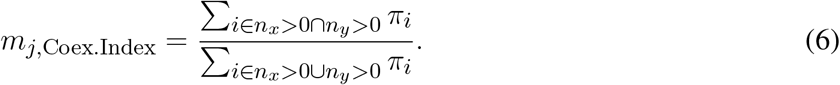

**Figure 4.**
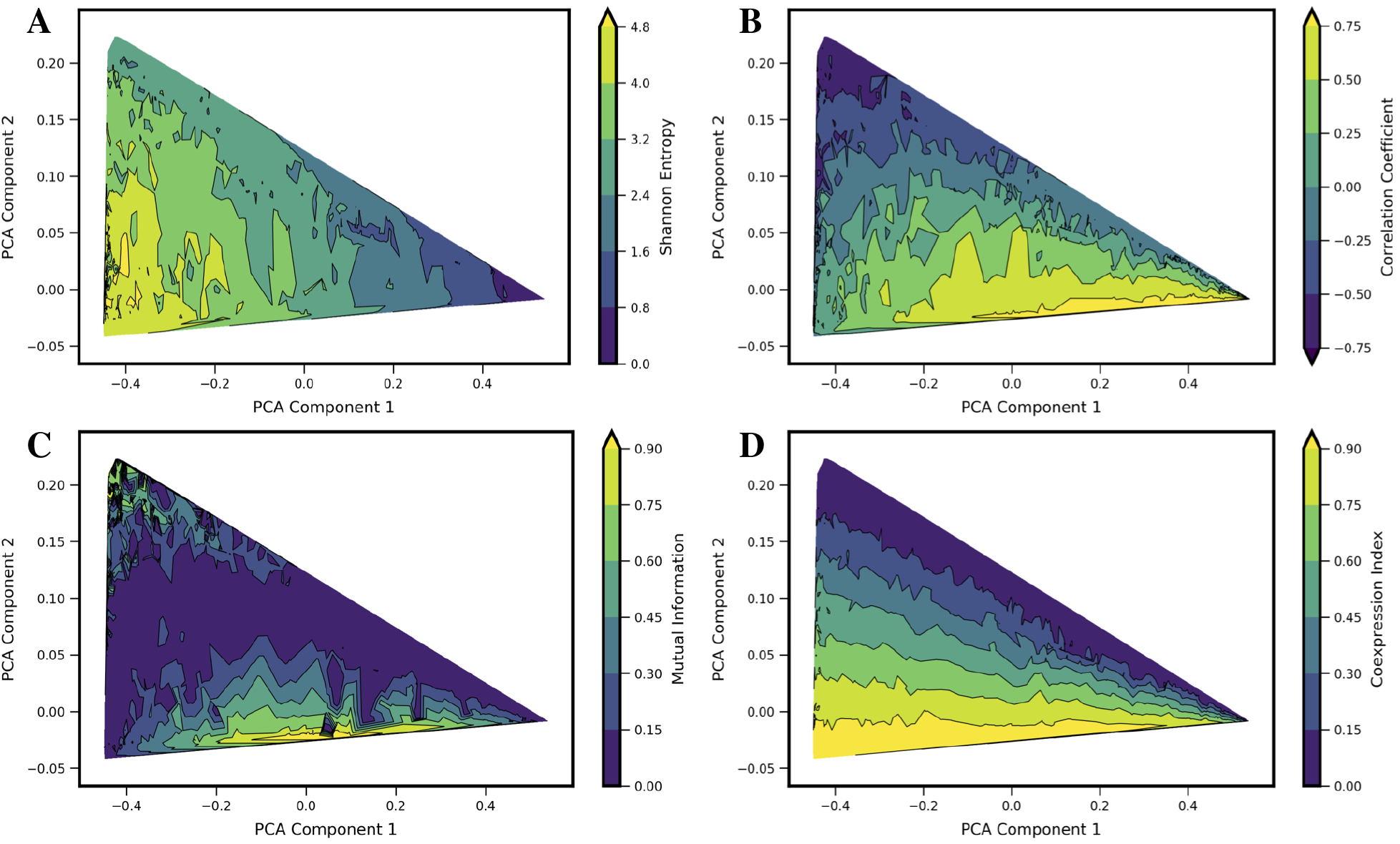
Comparison of four standard metrics of gene-gene coexpression with landscape shape. Metrics include: (**A**) Shannon Entropy. (**B**) Correlation Coefficient. (**C**) Mutual Information. (**D**) Coexpression Index (see text for details). Each metric was computed for each computed model landscape, using the same set of 34,097 Two-Gene Flex models as in Figs. 2 and 3. Contour plots show each metric as a function of principal components 1 and 2, obtained by local averaging and interpolation over the results from individual model landscapes. Taken together with Fig. 2, the results show how these metrics correspond with landscape shape.

We estimate the value of each metric as a function of landscape shape (that is, we estimate the function *m*(*c*_1_, *c*_2_), where *m* is a given metric and (*c*_1_, *c*_2_) are the coordinate values in PCA components 1 and 2). For each of the four metrics, we estimate and visualize this function by first computing each of the four metrics from the probability landscapes ***π***(*n*_*x*_, *n*_*y*_) corresponding to each of the 34,097 models. We then project the models onto the first two principal components, with a given metric serving as the colorscale (e.g., as shown with Shannon Entropy, Fig. 2 top). The continuous surface *m*(*c*_1_, *c*_2_) is then estimated by local averaging and interpolation over the computed results for each individual model landscape with the *tricontourf* routine from the matplotlib package. We found that each metric aligns in distinctive, and generally intuitive, ways with the PCA landscape shape-space. High or low values of each metric were to some extent localized to particular sub-regions of the triangle, and thus could be understood to be arising from landscapes of similar shape. However, numerous examples can also be found of models colocated (or nearly colocated) in the triangle but having different values of a given metric, so the functional dependence *m*(*c*_1_, *c*_2_) is noisy.

For Shannon entropy, the highest values are generally seen near the *hi/hi* vertex of the triangle, while the lowest values are seen near the *lo/lo* vertex. This reflects the amount of disorder in the *hi/hi* state of expression, in which a broad range of count-values are possible for each gene, whereas in the in the *lo/lo* vertex, count values are always zero or near-zero. The noise in expression levels can be quantified more precisely for the subset of models in the “slow-binding” regime (*h, f ≪ g, k*). In this parameter regime, cells show distinctive high (“*hi*”) and/or low(“*lo*”) expression states with mean counts *g*_*hi*_/*k* and *g*_*lo*_/*k*, respectively, and the disorder in each expression state can be quantified as Poisson birth/death noise (Al-Radhawi et al. (2019)), such that variance scales linearly with the expression rate *g*. Sources of disorder contributing to higher values of Shannon Entropy include both noisy expression within a given phenotype state and the ability for cells to exist in multiple different phenotype states (i.e., the breadth of a valley in the potential landscape, and the number of different valleys). Notably, in the parameter regimes studied here, the highest Shannon Entropy models are single-phenotype (*hi/hi*), indicating that the noise in this one state contributes more disorder than does noise from multiple phenotype-states. As such, models with two or more accessible states have intermediate values of Shannon Entropy.

A strongly negative correlation coefficient between the two genes is found near the *lo/hi*-*hi/lo* vertex of the triangle, which is occupied by models showing bistability (cells can express one gene or the other, but not both simultaneously) resulting from mutual repression in the network motif. Landscapes with high positive correlation tend to be those that combine expression in the *hi/hi* and *lo/lo* quadrants of the two dimensional subspace (see, e.g. 4B and 2D), resulting from mutual activation in the network motif. Mutual Information aligns somewhat with large absolute values of Correlation Coefficients, but cannot distinguish high positive from high negative correlation. Mutual Information values near zero co-localize with Correlation Coefficients near zero. This arc-shaped region bisecting the triangle also overlaps with the models lacking interactions between the two genes (see Fig. 3C).

The Coexpression Index shows the smoothest functional dependence on PCA components (*c*_1_, *c*_2_). Of note, the model-subspace of high coexpression is not fully overlapping with the subspace of high correlation coefficients. This reflects the fact that high simultaneous expression occurs in both genes in an uncorrelated manner, since the noise arises from aforementioned birth-death noise of mRNA transcription/degradation.

None of the four metrics are by themselves able to fully differentiate between landscape shapes. For example, model landscapes with similarly high values of Mutual Information include both *hi/lo*-*lo/hi* landscapes from mutual repression motifs and *hi/hi*-*lo/lo* landscapes from mutual activation motifs. (see, e.g., Fig. 4A and B). Model landscapes with similar intermediate values of Coexpression Index also encompass a variety of landscape shapes, including some that arise from different network motifs (see, e.g., Fig. 4C and D). Taken together, these results show that these four single metrics are not reliable determinants of landscape shape. They furthermore show that a given value for commonly used measures, as obtained from experimental data, can potentially arise from a variety of regulatory scenarios.

### 3.4 Stochastic theory-based analysis of coexpression landscapes from single-cell experiments reveals distinct developmental “landscape shape” trajectories

We applied the landscape shape analysis framework, developed above on the basis of theoretical models, to publicly available single cell RNA sequencing data in vertebrate development. We applied the analysis to putative MLP gene pairs in *Xenopus tropicalis* development collected at ten stages of embryonic development (Stages 8,10,11,12,13,14,16,18,20,22) (Briggs et al. (2018)). To carry out the analysis, we first analyzed the landscape shape-space for a restricted set of theoretical models, which encode only the MISA interaction motif. The MISA motif has been previously discovered to operate at critical cell-fate branch points (Graf and Enver (2009)) and has potential to enable both antagonistic expression and coexpression of genes in individual cells (depending on kinetic parameters), as is characteristic of MLP gene-pairs. We first generated a MISA-specific set of models for training the PCA shape analysis. In addition to restriction of the network motif, there were two other differences between the MISA-model training set (Fig. 5) and the Two-Gene Flex-model training set (Fig. 2). For MISA, we utilized quasipotential landscapes, rather than probability landscapes, in order to increase sensitivity to rarer cell-states (i.e., weaker landscape features). We furthermore restricted the kinetic parameters *h, f* to the fast (adiabatic) regime (see Table 1), in order to use the models to analyze time-resolved data. That is, the experiments measure embryos at different developmental stages, which are roughly 1-3 hours apart in time. We compare the steady-state landscapes from stochastic models to the experiment-derived landscapes at different timepoints by applying a quasi-steady-state assumption: we assume that the promoter-binding states (which govern gene activity) reach equilibrium faster than the progression of developmental stage, which is valid only in the adiabatic regime. Despite these modifications to the model training set, the projection of models onto the PCA subspace for MISA (Fig. 5) shows qualitative similarity to that of Two-Gene Flex ((Fig. 2), including delineation of a subregion of a triangle (note that the triangle is inverted between the two figures, which is an arbitrary result of eigenvector sign invariance). However, antagonistic expression of the two genes is a stronger feature across models in the MISA training set, such that the *hi/hi* vertex of the triangle for MISA still shows considerable probability for cells to antagonistically express one gene or the other (Fig. 5F).

**Figure 5.**
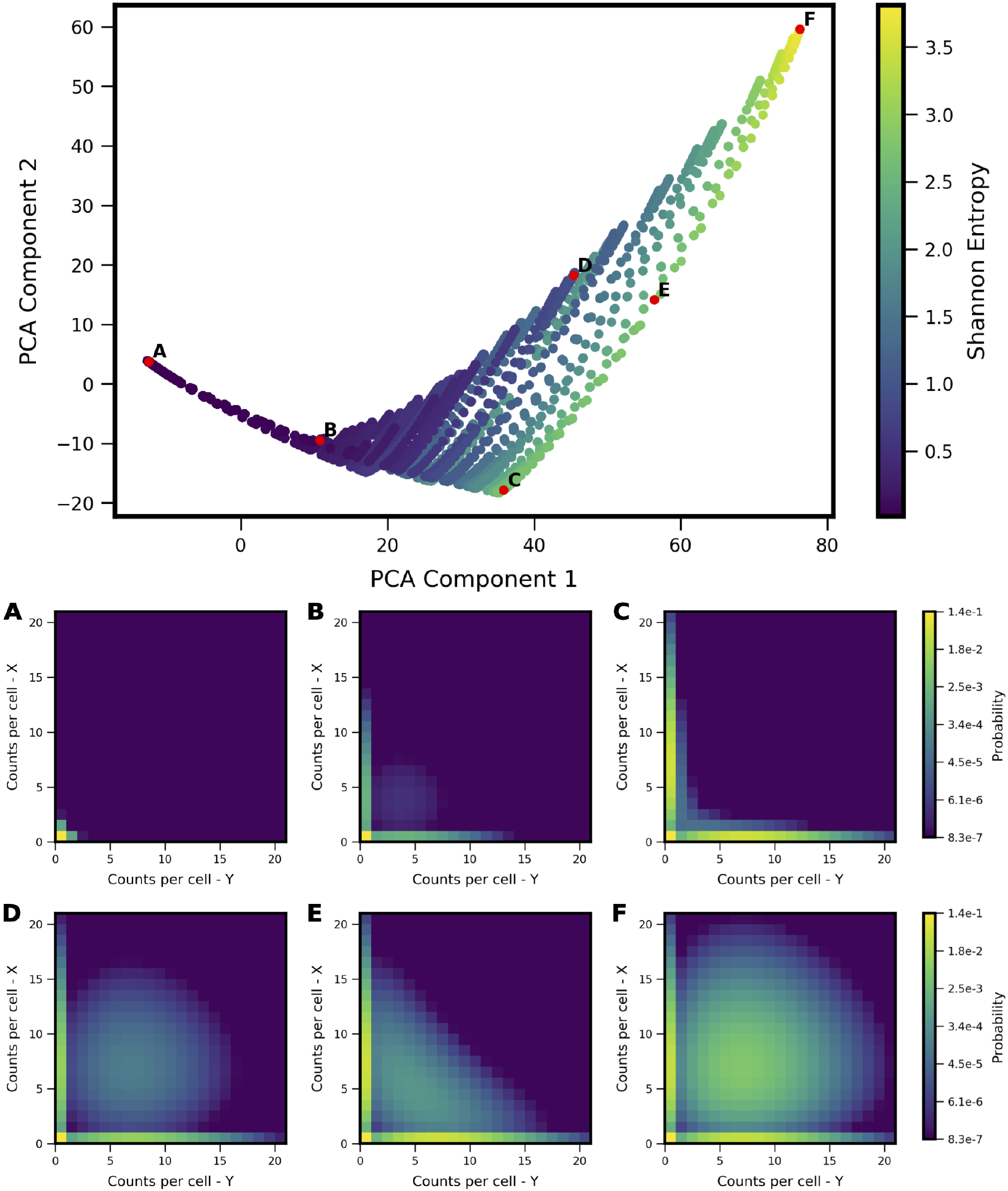
Shape-space of simulated MISA coexpression landscapes analyzed by PCA. Coexpression landscapes were computed for 22,718 unique two-gene stochastic network models with MISA logic and varying kinetic rate parameters. Promoter-state change rates were restricted to the fast regime (see Table 1). (Top) All model landscapes projected onto the first two principal components. Each dot corresponds to one model, colored by the model’s Shannon Entropy. (Bottom) Representative quasipotential landscapes *ϕ*(*n*) (see Text) of individual models from different regions of PCA component-space. Color of each discrete grid space in {*x, y*} corresponds to computed probability (in log-scale) to find a single cell with the corresponding numbers of {*x, y*} transcripts. (Analogous to Figure 2).

We extracted two-gene coexpression quasipotential landscapes corresponding to distinct developmental stages from the dataset of Briggs et al. We then projected the landscapes onto the PCA subspace, and thereby derived developmental trajectories through landscape shape-space. By way of illustration, we first present developmental trajectories for three representative gene pairs (Fig. 6). Gata5 and pax8 were identified (in Briggs et al.) as being antagonistically expressed within the intermediate mesoderm lineage, in cardiac mesoderm and pronephric mesenchyme cell subtypes, respectively. In contrast, lhx1 and pax8 were shown to coexpress in cells of the pronephric mesenchyme. Finally, the gene pair sox2 and brachyury (t) has been identified as influencing the cell fate decision between the neural plate and the dorsal marginal zone (Wardle and Smith (2004)), and was identified as presenting MLP behavior, characterized by high coexpression at some stage of development, followed by antagonistic expression at a later stage (Briggs et al. (2018)). We found that these three gene pairs showed distinctive trajectories through PCA subspace. All of the genes showed low expression early in development (stage 8) and their landscapes were colocated near the *lo/lo* vertex in the model subspace. Their trajectories then diverged: gata5-pax8 travels along the bistable edge of the triangle, increasing expression of both genes over the course of development, but in largely non-overlapping subpopulations of cells. In contrast, lhx1-pax8 shows strong coexpression starting at stage 14, and continues thereafter to move toward increasing values of PCA component 2, which coincides with increasing coexpression. (lhx1-pax8 landscapes for some of the measured developmental stages fall slightly outside the area reached by MISA models in the training set, suggesting that the interaction is likely not well described by a MISA motif). Finally, sox2-t shows a cyclic pattern in the shape subspace, where landscapes move towards hi/hi, and then back towards the antagonistic *lo/hi*-*hi/lo* region, landing in a similar area to gata5-pax8. Relating these landscape-shape dynamics to the stochastic MISA model parameters suggests that the gene-pairs undergo changes in the relative balance of mutual inhibition versus self-activation as development progresses (see Fig. S1).

**Figure 6.**
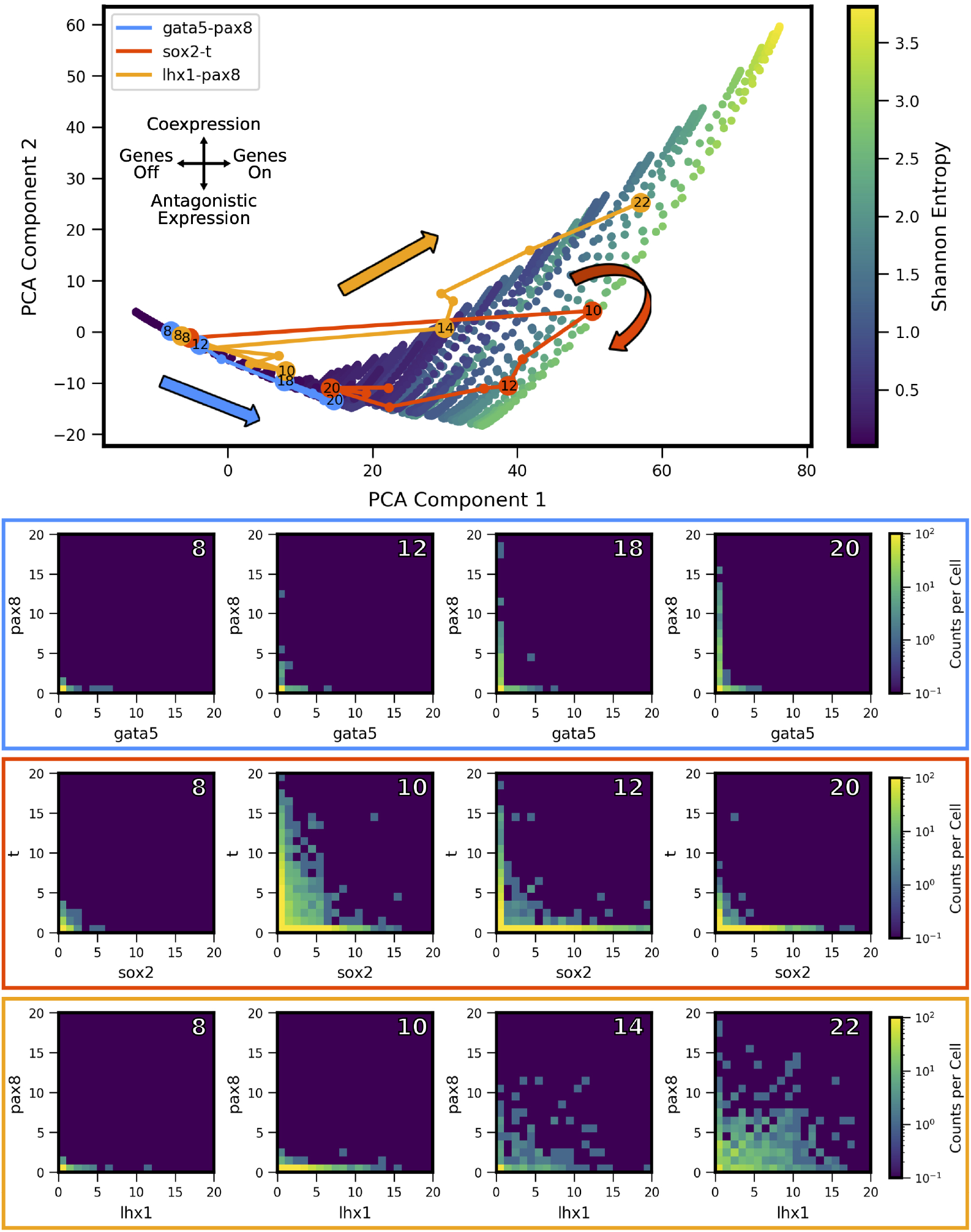
Landscape-shape trajectories of three representative gene pairs from scRNA-seq measurements in *Xenopus tropicalis* embryonic development. (Top) Developmental trajectories of three different gene pairs, plotted in principal component-space. Stages of interest shown below are labeled with the corresponding stage. Note the three stage 8 points are overlapping near the origin as a result of low expression. (Bottom) Coexpression quasipotential landscapes extracted from experimental measurements for the three gene pairs at different labeled stages of embryonic development (white numbers indicate developmental stage). The experiment-derived landscapes were trained on the principal components generated from the simulated MISA dataset of Fig. 5. Principal component 1 corresponds to overall level of expression, while component 2 separates antagonistic vs coexpression (see Fig. 7). The landscape of gata5-pax8 (blue) shows increasing antagonistic expression, consistent with movement along the lower left edge of the triangle in PCA shape-space. Sox2-t (red) shows high coexpression at stage 10, followed by later antagonistic expression, corresponding to a partial loop through PCA space, consistent with Multilineage Priming behavior. Lhx1-pax8 (orange) shows consistently increasing coexpression, corresponding to a mostly steady increase in principal components 1 and 2. (Data from Briggs et al. (2018)).

The experiment-derived developmental trajectories can be further understood by considering the features extracted by individual (by definition orthogonal) PCA components. Visualization of the first three PCA eigenvectors (Fig. 7) reveals that the first component (69.3% of covariance across the training set) can be summarized as separating landscapes with more or less expression overall, regardless of whether expression occurs in individual genes or both simultaneously. By contrast, the second component (15.6% of covariance) separates landscapes with coexpression versus antagonistic expression. The third component (6.8% of covariance) distinguishes landscapes with asymmetry between the two genes (subsequent components that describe less of the covariance displayed more complex shapes, and are not shown here). Comparison of the PCA scores versus developmental stage (Fig. 7, right) to the experiment-derived landscapes of Fig. 6 confirms visually that the PCA components extract the above-described features. For example, all three gene pairs show varying degrees of asymmetry (imbalance in expression levels of the two genes). Gata5-pax8 shows generally increasing positive amplitude of asymmetry, corresponding to stronger pax8 expression. At later stages, the other two gene-pairs show asymmetry in the other direction, corresponding to negative amplitude in component 3. Sox2-t exhibits a switch in asymmetry between stage 10 (t>sox2) and later stages (sox2>t).

**Figure 7.**
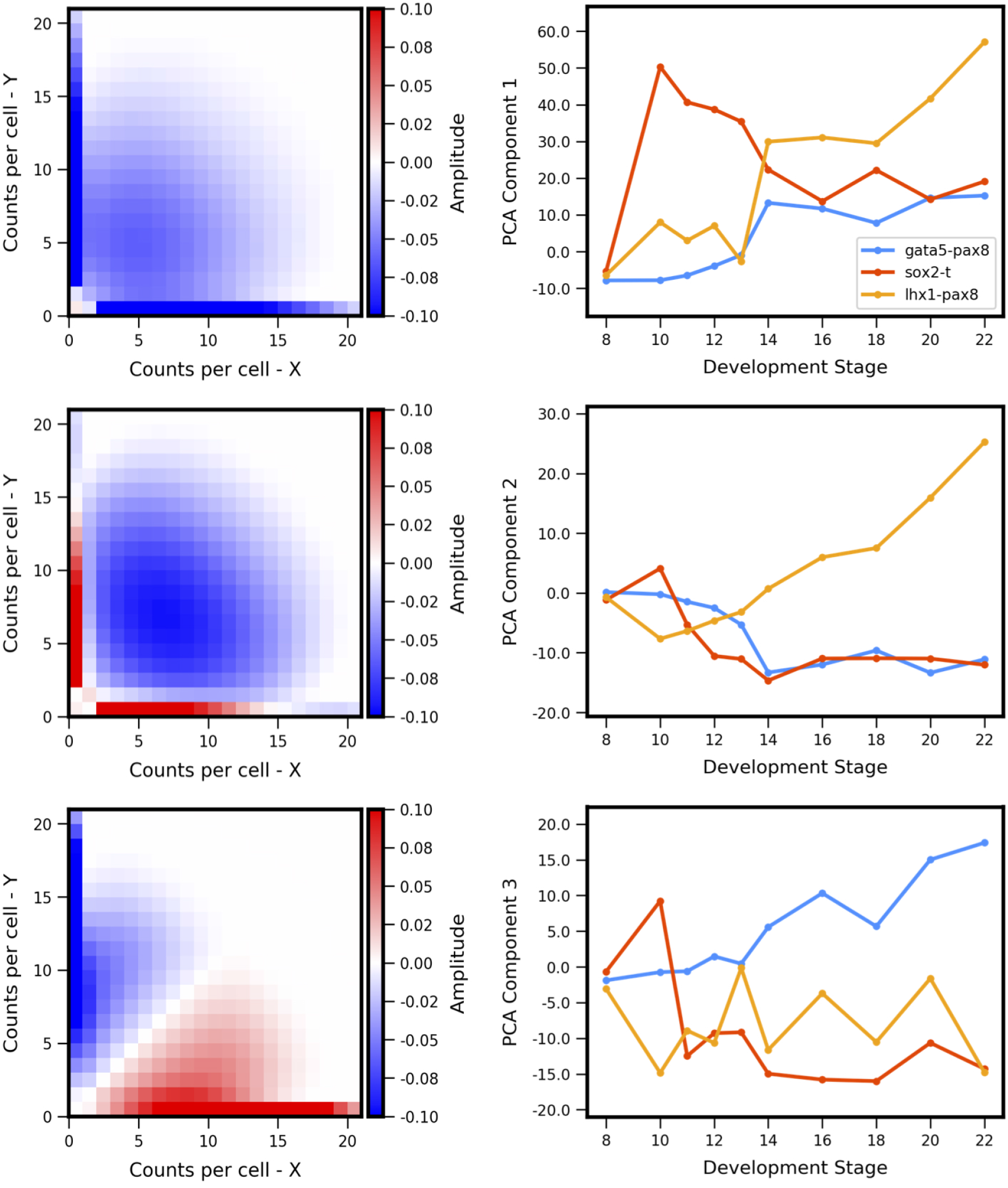
Principal components of landscape shape features. (**Left Column**) The reshaped PCA principal axes in feature space which represent the maximum variance in the data, specifically which features of the coexpression landscape that each component is accounting for. (**Right Column**) Magnitude or positive/negative value shift in observed variance for the respective component for each gene pair, versus developmental stage. Each component summarizes a landscape shape features: (**Top Row**) The overall amount of gene expression, (**Middle Row**) Antagonistic Expression vs Coexpression of the two genes, and (**Bottom Row**) degree of asymmetric expression between the two genes.

Developmental trajectories through the coexpression shape-space were compiled for 1,380 gene pairs (putative MLP pairs in *Xenopus tropicalis* identified by Briggs et al. (2018)). By applying the developmental trajectory clustering procedure described in Methods, we found that the trajectories of multiple gene pairs across different lineages display conserved patterns of coexpression dynamics. Twenty-four clusters were identified (see Supplemental Figs. S3 and S4), four of which are shown in Fig. 8; these clusters are chosen as representative of the different types of dynamic patterns obtained. The clusters display a variety of behaviors. For example, the cluster of Fig. 8B shows behavior that is consistent with MLP, i.e., genes are first increasingly coexpressed in single cells, followed by a switch towards antagonistic expression, similar to the cycle in PCA space delineated by sox2-t in Fig. 6. Surprisingly, we also observed clusters that show “inverted MLP” behavior (Fig. 8A) where the genes initially turn on in non-overlapping subsets of cells (i.e., increasing antagonism), but later show increasing coexpression in single cells. A number of the analyzed gene pairs showed generally antagonistic expression (Fig. 8C), reminiscent of gata5-pax8. Others showed behavior consistent with the dynamics of MLP (i.e., first coexpression, later antagonistic expression), but with coexpression being only weakly detectable (Fig. 8D). The gene pairs represented in these clusters include (but are not limited to) regulators of embryonic development including zic3, hoxc10, and neurog1. The full list of clusters and their associated gene pairs are listed in the Supplementary File 1.

**Figure 8.**
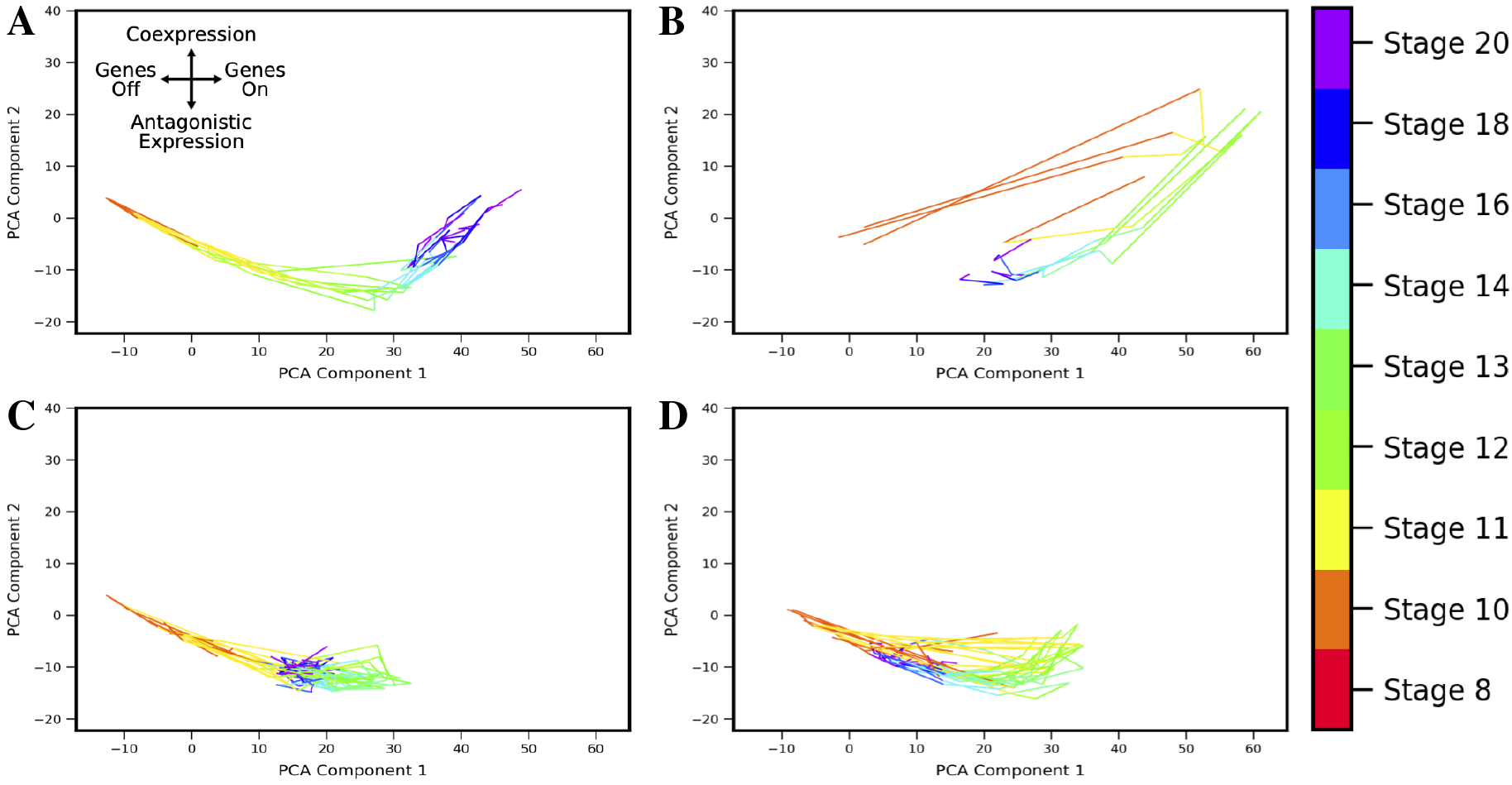
Landscape shape trajectory clustering reveals conserved patterns of gene-pair coexpression dynamics during development. Four representative trajectory clusters showing distinct dynamics are presented (full list of 24 clusters and associated gene pairs in Supplement). Gene pairs in cluster **A** display behavior of an “inverted MLP”: first undergoing increasing antagonistic expression which then switches to increasing coexpression around stage 13. Gene pairs in cluster **B** follow the typical MLP behavior, with highest coexpression taking place around stage 10 followed by antagonistic expression at later stages. Cluster **C** shows consistent antagonistic expression (negative component 2), with nonmonotonic overall expression (a switch-back in component 1 around stage 12). **D** shows cyclic behavior similar to **B**, with highest coexpression at stage 12, but overall expression and relative amount of coexpression is lower.

## 4 DISCUSSION

In this work, we comprehensively studied theoretically predicted single-cell gene-gene coexpression landscapes based on a class of stochastic gene regulation models, and applied the theory to analyze two-gene coexpression landscapes from single cell measurements. From a training set of tens of thousands of computed, theoretical landscapes, we identify Principal Components of landscape covariance that serve as simple “fingerprints” of landscape shape and reflect underlying gene-gene interaction dynamics. We then apply the theoretically-derived framework to scRNA-seq data from vertebrate development. In so doing, we uncover distinctive and novel developmental trajectories of gene-gene coexpression. Specifically, our framework reveals a nuanced picture of multilineage priming, where the relative balance between expression of gene pairs simultaneously (in the same cells) versus antagonistically (in different cells) within a lineage shows complex dynamics during development, for example, revealing that simultaneous coexpression occurs either earlier or later than antagonism. Based on the results, we propose that the framework developed here can be generalized to other single cell datasets and stochastic network models to analyze the evolution of gene-gene regulatory interactions over the course of development.

The theoretical framework applied here–discrete, stochastic reaction kinetic modeling–is well-suited to aid interpretation of single cell measurements: first, because it inherently captures cell population heterogeneity and second, because of the direct correspondence between the computed quantities (e.g., probability to find a given number of mRNAs in a cell) and experimentally-measured transcript counts in scRNA-seq. The theoretical models can partially reproduce true cell population heterogeneity, but also neglect many sources of noise, both biological and technical. We employ models that treat intrinsic noise but neglect sources of persistent cell-to-cell variability (i.e., extrinsic noise) (Swain et al. (2002)), which is known to contribute to noise in gene expression. For example, one source of extrinsic noise would be asynchronicity between cells, where individual cells might be at different stages of progression in development. Here, we opted to use a relatively simplistic model framework (i.e., no additional noise assumptions beyond intrinsic noise of biomolecular interactions, relatively few reactions describing molecular mechanisms of gene regulation, etc.) to minimize the number of model parameters while still enabling study of a variety of “rules” for gene regulatory logic. The framework presented here could be expanded in the future by integration of additional types of mechanistic assumptions and noise sources in the stochastic models.

The models also neglect technical noise/measurement errors arising from experiments (Grün et al. (2014)). For example, scRNA-seq measurements face a well-known technical issue of drop-outs (Kharchenko et al. (2014)), which we have not included in our modeling. Future efforts may improve the presented modeling framework by inclusion of these additional sources of noise, or by additional data-processing steps for imputation of missing datapoints (Gong et al. (2018)). However, such an approach would also present challenges by necessarily introducing additional assumptions about cell population heterogeneity, which is still not fully understood. Given the danger of false signals (Andrews and Hemberg (2019)), we opted here to utilize minimal data processing in comparing our theoretical results to a public dataset. We also note that the discrete stochastic modeling framework advanced in this work has potential to shed new light on the drop-outs issue: a relatively large proportion of “zeros” arises naturally from discrete stochastic models, depending on the regulatory interactions among genes, suggesting that perhaps biological variability plays a larger role in producing dropouts than has previously been supposed. Overall, despite the lack of additional biological/technical noise sources in our models, we note that our computed landscapes qualitatively reproduce the noise characteristics of the scRNA-seq measurements, in that they showed similarly broad distributions of coexpression. Thus we conclude that the simplistic models employed here are sufficient for the current application, which focused on characterization of coexpression landscape shape and its evolution in development, but we also foresee that incorporation of additional noise sources in the model might improve the practical utility of our proposed coexpression-shape-based analysis.

We focused here on two-gene models and pairwise interactions, because (1) certain gene-pairs are known to play a critical role in development (Graf and Enver (2009)) (2) the edges (pairwise interactions) are the elemental units or building blocks of larger regulatory networks. However, the focus on pairwise interactions has potential drawbacks: it does not elucidate how gene-pair interactions are modified when embedded in a larger network. In the same vein, it does not differentiate between direct or indirect interactions between genes (e.g., by direct transcriptional regulation versus molecular intermediaries). In principle, the framework presented here could be expanded to treat “3-body” (or higher order) interactions among genes, though this presents several computational challenges. For example, solution of the CME becomes intractable already for 3-gene networks, such that advanced approximation methods (Zhang and Wolynes (2014)) or more costly simulations (Tse et al. (2018)) become necessary. Nevertheless, expansion of the approach to higher-order interactions is feasible, and recent work has revealed how such as approach might proceed, for example, by incorporating developments in multivariate information measures (Chan et al. (2017)).

In this work, linear PCA was used to identify shape features of gene-pair coexpression landscapes, and this approach was useful for separating landscapes with, e.g., more simultaneous coexpression versus more antagonistic expression for a given gene pair. Another possible extension of the method in the future could be to test alternative, nonlinear dimensionality reduction strategies for potential improvements in classifying coexpression landscapes based on desired features.

## Supporting information

Supplemental Section

## AUTHOR CONTRIBUTIONS

Conceptualization and study design, CG and ER; Coding, CG; Analysis, CG, HR, and ER; Visualization, CG and HR; Writing, review, and editing, CG, HR, ER. All authors read and approved the final version of the manuscript.

## SUPPLEMENTAL DATA

The Supplementary Material for this article can be found online.

## DATA AVAILABILITY STATEMENT

The simulation codes produced for this study can be found at https://github.com/Read-Lab-UCI/Stochastic_Coexpression_Landscape_Analysis.

