## Supplemental Section for "Analysis of single-cell gene pair coexpression landscapes by stochastic kinetic modeling reveals gene-pair interactions in development"

### Supplemental Information

#### 1 Supplemental Methods: Stochastic Gene Network Models

##### 1.1 Model Reactions

The full reactions of the biochemical network are as follows.

Promoter state change reactions (followed by [forward rate, reverse rate]):

| | |
|---|---|
| $X_{00} \rightleftharpoons X_{01}$ | $h_a x^2/2, f_a$ |
| $X_{00} \rightleftharpoons X_{10}$ | $h_r y^2/2, f_r$ |
| $X_{01} \rightleftharpoons X_{11}$ | $h_r y^2/2, f_r$ |
| $X_{10} \rightleftharpoons X_{11}$ | $h_a x^2/2, f_a$ |
| $Y_{00} \rightleftharpoons Y_{01}$ | $h_a y^2/2, f_a$ |
| $Y_{00} \rightleftharpoons Y_{10}$ | $h_r x^2/2, f_r$ |
| $Y_{01} \rightleftharpoons Y_{11}$ | $h_r x^2/2, f_r$ |
| $Y_{10} \rightleftharpoons Y_{11}$ | $h_a y^2/2, f_a$ |

mRNA synthesis reactions:

$$X_{00} \xrightarrow{{g_{00}}^X} X_{00} + x$$
$$X_{01} \xrightarrow{{g_{01}}^X} X_{01} + x$$
$$X_{10} \xrightarrow{{g_{10}}^X} X_{10} + x$$
$$X_{11} \xrightarrow{{g_{11}}^X} X_{11} + x$$
$$Y_{00} \xrightarrow{{g_{00}}^Y} Y_{00} + y$$
$$Y_{01} \xrightarrow{{g_{01}}^Y} Y_{01} + y$$
$$Y_{10} \xrightarrow{{g_{10}}^Y} Y_{10} + y$$
$$Y_{11} \xrightarrow{{g_{11}}^Y} Y_{11} + y$$

mRNA degradation reactions:

$$x \xrightarrow{k} 0$$
$$y \xrightarrow{k} 0$$

Our reaction notation and formalism are based on previous work ( [3]). We label two genes $X$ and $Y$. Each gene encodes a protein, which acts as a transcription factor that potentially regulates its own expression as well as that of the other gene. Each gene has a promoter (or more generally, regulatory regions of DNA) that can be bound by any combination of its own expressed protein and/or the other gene's expressed protein. The promoter states are thus labeled as: $X_{00}$ (neither transcription factor is bound to X's promoter), $X_{01}$ (X's own protein is bound, resulting in auto-regulation of gene expression), $X_{10}$ (Y's protein is bound to X's promoter, resulting in cross-regulation), $X_{11}$ (both proteins are bound to X's promoter, resulting in combinatorial regulation). (The promoter states for gene $Y$ are defined in the same manner.) The regulatory effect of each promoter state (i.e., the effect of having none, one, or both proteins bound on the gene's expression) is accounted for by the transcription rates: e.g., when gene $X$'s promoter is unbound, it transcribes at rate $g_{00}^X$. Binding of $Y$'s protein $X$ changes the transcription rate to $g_{10}^X$, which may be lower, higher, or the same, depending on whether the effect of $Y$ on $X$ is assumed to be repressing, activating, or not impacting. All other transcription rates for each promoter state and for gene $Y$ are defined similarly. The model describes mRNA numbers in the cell, to enable direct comparison with single cell transcriptomic data. Translation of mRNA into protein is not explicitly accounted for in the model. mRNA molecules are denoted with lower-case letters (e.g., mRNA from gene $Y$ is denoted $y$). mRNAs are degraded at rate $k$.

The changes of promoter state occur with rate constants denoted $h$ (when the change of state occurs due to a transcription factor binding event) and $f$ (when the change of state occurs due to an unbinding event). We make the common assumption that homodimeric transcription factors regulate gene expression, which is a general and convenient way to include cooperativity in the model. Since protein molecules are not explicitly included in the model, the rates of promoter-state-change due to protein binding to DNA are expressed in terms of mRNA number. For example, the change of promoter state $X_{00} \rightarrow X_{01}$ occurs with rate $hx^2/2$, where $h$ is a rate parameter and $x$ is the copy number of mRNAs encoded by gene $X$ (details on the parameter $h$ and justification for the quadratic dependence on mRNA is are given in the next section). Promoter-state-changes that occur due to transcription factor unbinding (rate $f$) are considered first-order reactions that do not depend on the concentration of free protein (or mRNA). The rates are further denoted by subscripts: $h_a$ and $f_a$ are rates of change due to binding and unbinding, respectively, of the auto-regulator (i.e., they involve the self-protein) and $h_r$ and $f_r$ are rates of change due to binding and unbinding, respectively, of the cross-regulator.

#### 1.2 Model Parameters

The models employed in this work are intended to be phenomenological, nevertheless we assign rate constants to intracellular processes that are in line with experimental estimates from vertebrates, where possible. In particular, the fast rates of promoter state change from unbinding ($f$) or binding ($h$) are derived from dissociation and association rate constants reported in [2], and we assume first-order kinetics of unbinding. They report values of transcription factor dissociation rates in the range $10^{-3}$ to 10 per second, which we convert to the range 10 to $10^5$ per hour.

Here, we discuss in more detail the promoter-state-change due to binding reactions. Consider the promoter-state-change:

$$\mathrm{X_{state1}} \longrightarrow \mathrm{X_{state2}}$$

According to standard biochemical kinetics, if binding of factor F causes the state-change, the rate of this reaction is given by

$$b\mathrm{F}$$

where $b$ is the binding rate of the transcription factor to DNA (units $\mathrm{M}^{-1} \cdot \mathrm{s}^{-1}$) and F is the concentration of the transcription factor in the nucleus (units M) that effects the change of state. Assuming the transcription factor is a homodimer, with the corresponding monomer labeled R, then invoking a quasi-steady-state approximation according to standard kinetic rate laws, the dimer can be expressed in terms of the monomer by

$$\mathrm{F} = \frac{\mathrm{R}^2}{2K_d},$$

where $K_d$ is the equilibrium dissociation constant for the dimerization reaction. If we further assume that R is linearly proportional to its corresponding mRNA $m$ with proportionality factor $\gamma$ ($R = \gamma m$), then the rate of promoter-state-change due to binding is given by:

$$b\mathrm{F} = b\frac{(\gamma m)^2}{2K_d} = hm^2/2.$$

Thus, under these assumptions the promoter-state-change rate has quadratic dependence on mRNA number, and our parameter $h$ relates to the transcription factor binding rate, the ratio of protein to mRNA, and the equilibrium binding affinity of the homodimeric transcription factor.

According to the transcription factor binding rates reported in [2], in the range $10^6$ to $10^8$ $M^{-1}s^{-1}$, and assuming a nuclear concentration of transcription factor of $F = 10$ nM, the promoter-state-change-due-to-binding occurs with rates ranging from $bF$ =100 to 4000 /hour. To achieve comparable rates, given average mRNA counts of approximately 4 per cell in our model (in on-state), we estimate $h \approx 10 - 500$.

Note that promoter-state-changes can occur not only due to simple binding events, but also due to more complex, multi-step processes involving binding/unbinding, cooperative molecular interactions, and chromatin structural rearrangements. These processes are expected to be generally much slower than binding and unbinding events, but may still occur with rates that are functionally dependent on the amount of some essential molecular regulator present in the nucleus. In order to explore the possibility of these types of slow regulatory changes due to chromatin structural rearrangements, we also explore parameter values of $h$ and $f$ that are orders of magnitude slower than these estimates based on transcription factor binding/unbinding (see Main text and Table 1).

#### 1.3 Master Equation for Gene Regulatory Network Model

The Master Equation describes the time evolution of the probability of the system to be in any given state. Following Zhang and Wolynes ( [3]), we can express the stochastic expression of one gene by

$$\mathbf{P}(n,t) = \begin{bmatrix} P_{00}(n,t) \\ P_{01}(n,t) \\ P_{10}(n,t) \\ P_{11}(n,t) \end{bmatrix}$$

where, e.g., $P_{00}(n,t)$ is the probability of having n copies of mRNA, given that the promoter is in state $\{00\}$, at time $t$ (note that our model definitions differ slightly from those of Zhang and Wolynes, namely in that we model mRNA rather than protein. However, the mathematical formalism is identical). Then the Master Equation for this particular gene can be expressed as:

$$\begin{aligned} \frac{d}{dt}\mathbf{P}(n,t) = \mathbf{g}\{\mathbf{P}(n-1,t) - \mathbf{P}(n,t)\} \\ +\mathbf{k}\{(n+1)\mathbf{P}(n+1,t) - n\mathbf{P}(n,t)\} \\ +\mathbf{w}\mathbf{P}(n,t) \end{aligned} \tag{1}$$

Here, the first term accounts for transcription and $\mathbf{g}$ is a diagonal $4 \times 4$ matrix with transcription rates as diagonal elements $\{g_{00}, g_{01}, g_{10}, g_{11}\}$ (which may be different depending on whether gene $X$ or gene $Y$ is considered). The second term describes mRNA degradation, and $\mathbf{k}$ is a diagonal matrix with diagonal elements all equal to $k$ (since the promoter state is not assumed to affect the mRNA degradation rate). The last term accounts for the rate of transitions between promoter states, e.g., in the matrix $\mathbf{w}$, using the above ordering of promoter states, the element $w_{21}$ holds the transition from state index 1 ($\{00\}$) to state index 2 ($\{01\}$) and is thus given by the rate $h_a n^2/2$. If the direct promoter-state-change from state $i$ to state $j$ is not possible (e.g., $\{00\}$ cannot directly change to $\{11\}$ in one reaction event) then the element $w_{ji} = 0$. Also, for probability conservation $w_{ii} = -\sum_{j \neq i} w_{ji}$.

The Master Equation (1) is incomplete, since it only treats one gene at a time. Eqn 1 is expanded to simultaneously account for transcript levels and promoter states of both genes $X$ and $Y$. The instantaneous state of the system is given by vector $\mathbf{n} = [n_X, n_Y, X_{ij}, Y_{ij}]$ where here $i, j \in \{0, x, y\}$ (i.e., the system is fully described by enumerating the mRNA copy numbers and identifying the promoter states for both genes). If both types of mRNA are assumed to exist in the cell in copy numbers that never exceed $M - 1$, then the total size of the enumerated space including all possible states is $N = M \times M \times 4 \times 4$ (note that the total number of protein copy number states includes the state of 0 copies, thus the $M$ copy-number states are $\{0, 1, ...M - 1\}$. Thus, in order to describe the complete time-evolution of the system, simultaneously accounting for both genes and their auto- and cross-regulatory effects, the probability is described by a time-varying state vector $\mathbf{P}(\mathbf{n}, t)$ of length $N$. Furthermore, the full system Master Equation can be constructed (expanding on the above formalism) in vector-matrix form with reaction rate matrices of size $N \times N$. For probability conservation, $\sum_i^N \mathbf{P}(\mathbf{n}_i, t) = 1$.

### 2 Supplemental Figures

#### 2.1 Effect of binding parameters

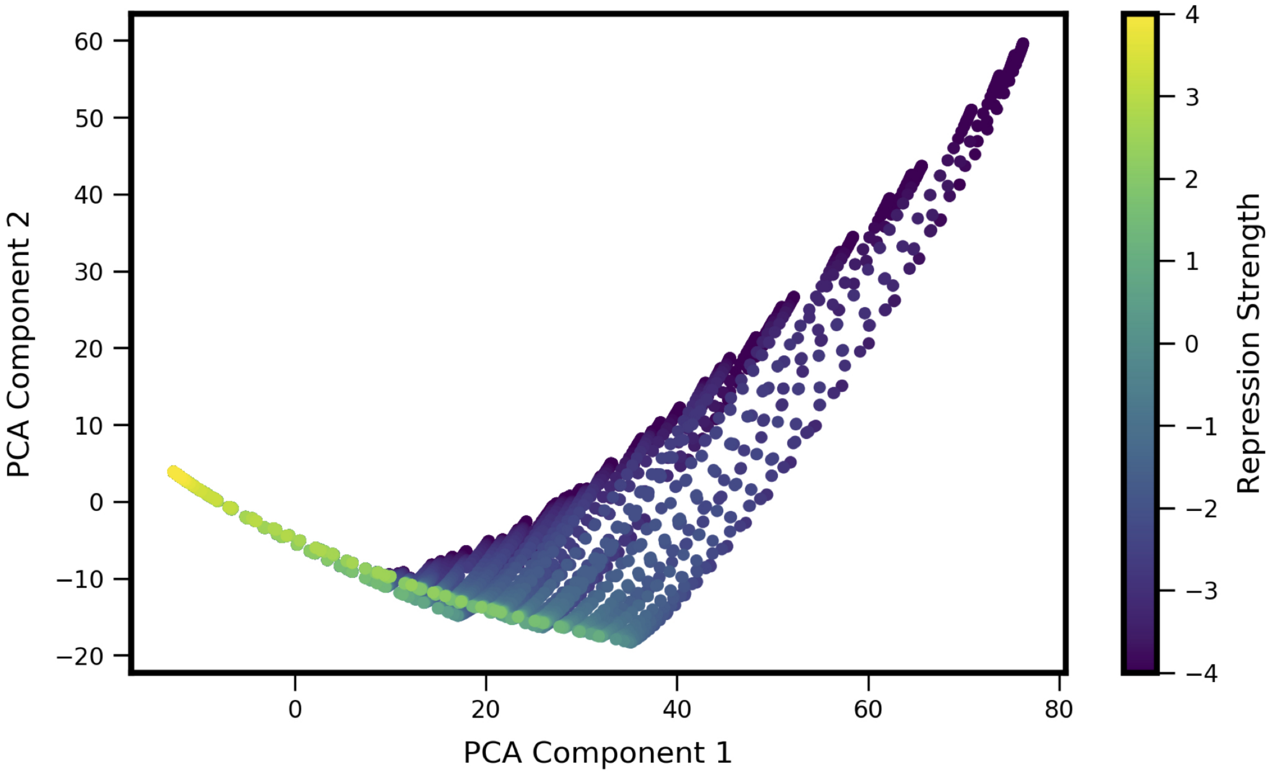


Figure S1: Relationship between kinetic parameters (binding/unbinding) and coexpression landscape shape for the MISA model. Repression strength is defined as $log(X_a/X_r)$, where $X_a = f_a/h_a$ is a dissociation constant for activator binding (i.e., the ratio of off-rate to on-rate of the activator (self-TF)) and $X_r = f_r/h_r$ is the corresponding dissociation constant for repressor binding (the other TF)). The highest values of coexpression (top-right) correspond to lowest repression strength, i.e., self-activation outweighs mutual inhibition.

#### 2.2 State Space Truncation Approximation Error

In the Main Text figures, we assume that mRNA copy numbers never exceed $M - 1 = 20$. This is an approximation that neglects the low probability of mRNA numbers exceeding $M - 1$, though in principle mRNA copies can take on any non-negative integer value. To check the validity of this approximation, we solved the steady-state CME for larger values of $M$ and computed the fraction of probability in the steady-state landscape present in the expanded (i.e., previously neglected) set of states. For a representative model with high coexpression, the total gained probability as a function of increasing $M$ from $21 \rightarrow 26$, $26 \rightarrow 31$, and $31 \rightarrow 36$ was 1.47e-5, 2.90e-8, and 2.27e-11 respectively. (Corresponding values were essentially 0 for models with low overall expression). This indicates that there is minimal gain to be had from increasing the state space beyond the original cutoff of $M$=21, supporting validity of the approximation. We furthermore found that the PCA shape-space over all models and resultant experimental trajectories were unchanged with an expanded state-space (Fig. S2).

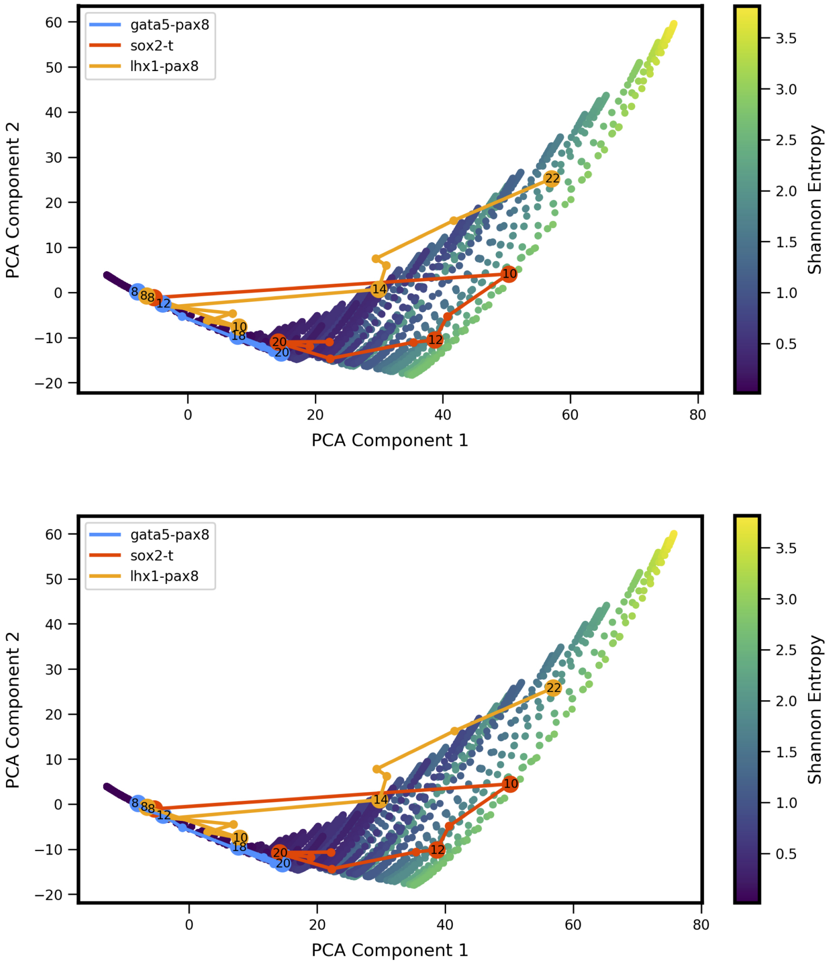


Figure S2: Comparison of $M$=21 (Top) to $M$=36 (Bottom) simulated state space. An expanded state space for the simulation training data results in a negligible difference in the PCA landscape and its application to the gene pair trajectories.

#### 2.3 Co-expression Trajectory Clustering

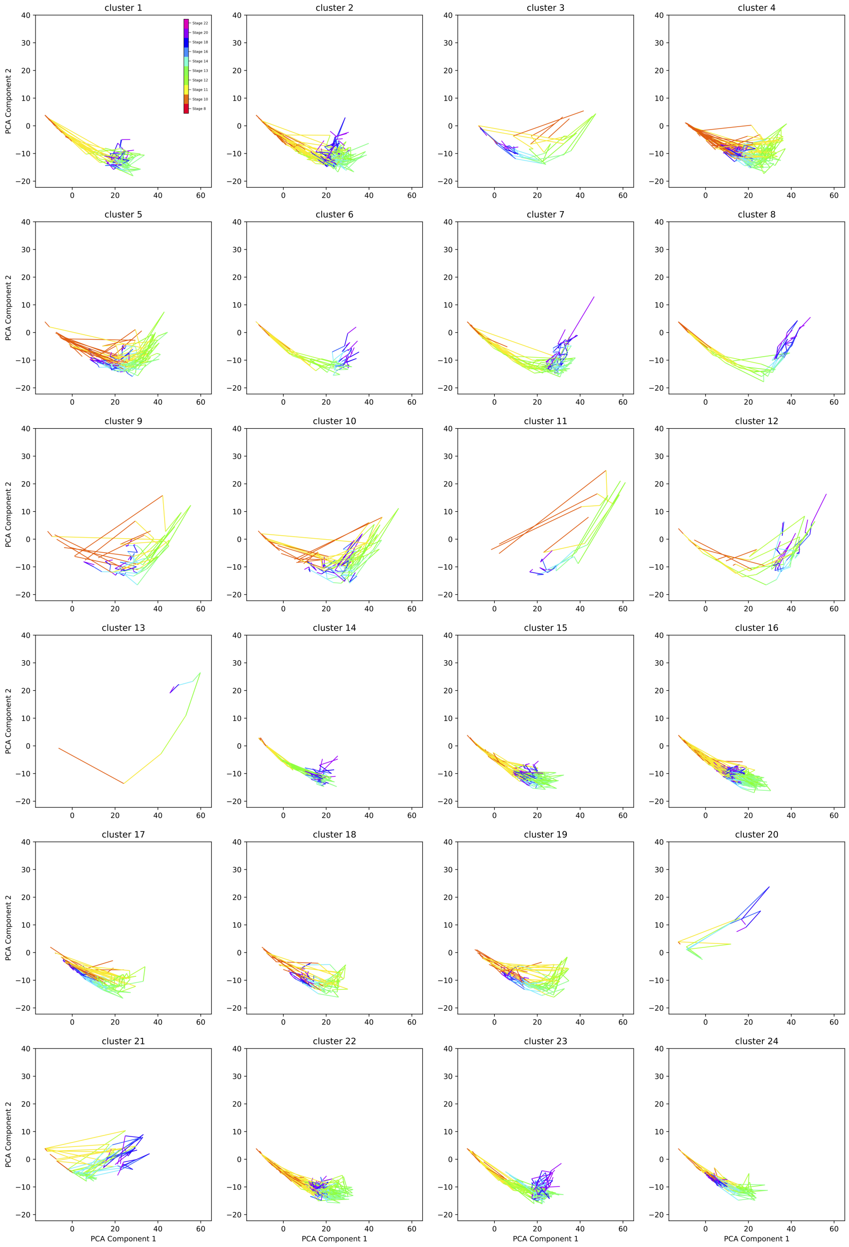


Figure S3: Developmental trajectory clustering results for the gene pairs proposed by [1]. Only clusters where > 50% of the trajectories have at least one stage where PCA component 1 is greater than 20 are shown, where PCA component 1 indicates the degree of overall expression. This filtering method thus retains only landscape trajectories that show high expression during at least one developmental stage, since it was difficult to discern shape trajectories for gene pairs with low overall expression. This step retained 398 gene pairs out of 1380.

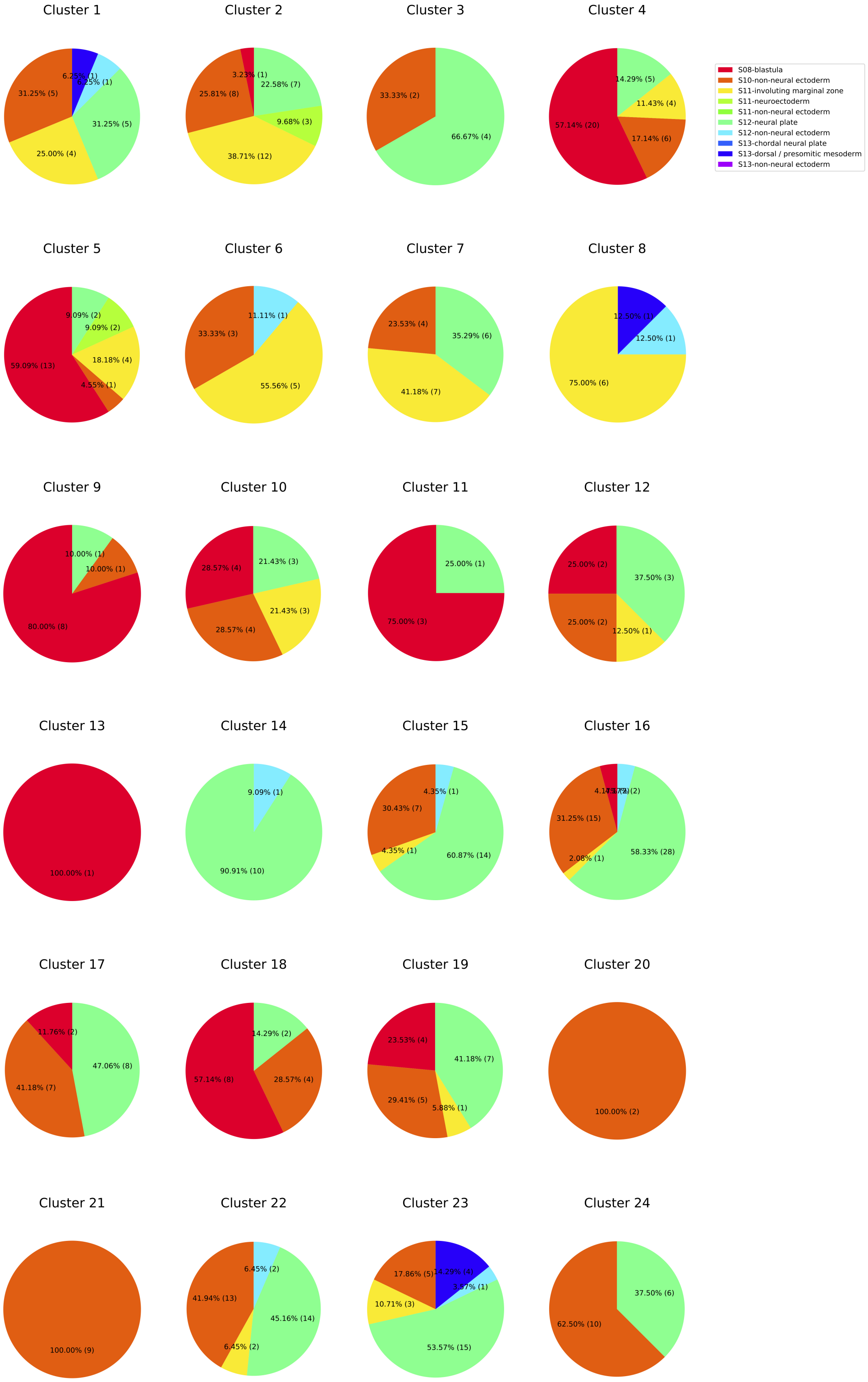


Figure S4: Pie chart of the root lineage branching point for the gene pairs in each cluster of Figure S3.

#### 2.4 Data-driven PCA Training

In this work, we use theory-driven analysis of gene-pair landscape shape. Alternatively, one can obtain a PCA shape-space directly from experimental data. In both cases (theory-driven versus data-driven), a low-dimensional description of landscape shapes is obtained. We analyzed the PCA shape-space generated directly from the same set of experimental scRNAseq landscapes from Briggs, et al. As shown in Fig. S5, in this data-driven space the three representative gene pairs show distinctive trajectories over the course of successive developmental stages, similar to the theory-driven results presented in Main Text. However, the choice of PCA training set influences the covariances and thus influences the shape features that are identified by each Principal Component. For example in Fig. S5, the second component is identifying asymmetry in the landscape, whereas in our MISA-derived theory-driven shape-space (Fig. 7), the second component identified simulaneous coexpression (versus mutually exclusive expression) while the third component identified asymmetry.

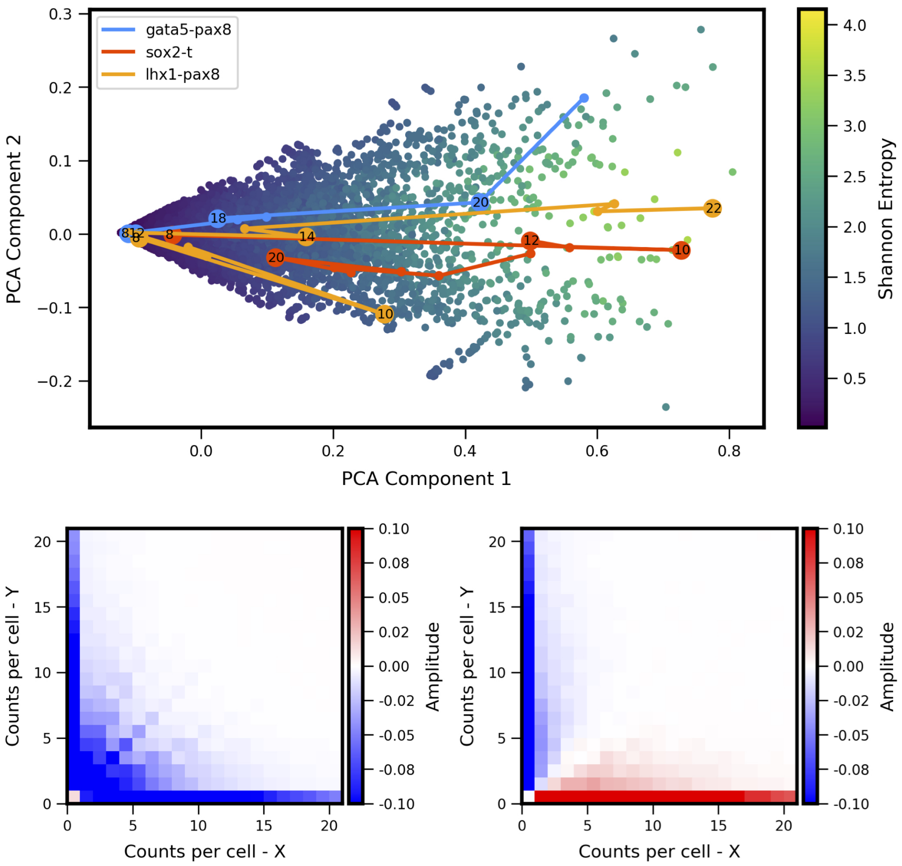


Figure S5: Data-driven PCA shape-space: (top) 13800 experiment-derived quasipotential landscapes (1380 gene pairs times 10 stages from Briggs, et al., as described in Main Text.) projected onto the first two Principal Components. Figure is analagous to Main Text Fig. 6 (top), showing developmental shape-space trajectories of the same three representative gene pairs (bottom) Eigenvectors corresponding to the first two components (analagous to Main Text Fig. 7).

### 3 Supplemental Data

Supplemental File 1: Cluster_gene_pair_members.csv

- **Gene pairs within the clustered trajectories**. Table with 398 rows by 3 columns, provided as .txt file with comma-delimited values. First row is column headers. Column 1: Cluster ID number the row is describing, corresponding to 24 trajectory clusters shown in S2. Columns 2 and 3: First and second gene symbols, respectively, analyzed to create the developmental trajectory.
